# Decoding the Circulating Proteome: Matrix and Immune Context Markers Shape Early Multi-Cancer Detection

**DOI:** 10.1101/2025.11.10.687473

**Authors:** Vivek Singh, Rashmi Kushwaha

**Affiliations:** Department of Biochemistry, King George’s Medical University, Lucknow, Uttar Pradesh, India 226003; Department of Pathology, King George’s Medical University, Lucknow, Uttar Pradesh, India 226003

**Keywords:** Multi-cancer early detection (MCED), Circulating proteome, Cross-platform validation, Matrix–immune–secretory axis, Liquid biopsy

## Abstract

**Background:** Blood-based proteomics offers a complementary path to multi-cancer early detection (MCED) by capturing the tumor secretome and host response. We analyze recent (2020–2025) evidence and add pathway/hallmark context, cross-platform validation, and proteome-scale protein–protein interaction (PPI) inference to guide translational panel design.

**Methods:** We reviewed extensive prospective and multi-cancer studies using Olink, SomaScan, and mass spectrometry, contrasting case-control versus prospective performance. Candidates were organized into Known and Novel sets and mapped with GeneCodis (GO/KEGG/Reactome) and Cancer Hallmarks. Clinical relevance was assessed using GEPIA 3.0 Cox Forest plots and TCGA-survival Kaplan–Meier curves (median split; log-rank). Protein-level corroboration used TPCPA/RPPA Z-score distributions across tumor types. To contextualize molecular crosstalk, we incorporated in silico PPI prediction to evaluate whether candidates cluster into interaction sub-modules relevant to secretory/TGF-β, matrix remodeling, and immune-follicular biology.

**Results:** Beyond classic antigens/inflammatory markers (e.g., CEACAM5, WFDC2/HE4, GDF15), the Novel set converged on a matrix-immune-secretory axis comprising ECM proteases (MMP12, ADAM8), antigen presentation/B-cell programs (CD74, CXCL13), secretory/TGF-β signaling (TGFB1), and epithelial invasion (CDCP1). Forest plots show adverse hazards for secretory/matrix genes across multiple epithelial cancers, while immune-follicular genes exhibited context-dependent effects; TCGA-survival curves reproduced these directions. TPCPA demonstrated tumor-type–specific protein elevation, supporting detectability. PPI inference organized candidates into coherent interaction modules (e.g., TGFB1, CDCP1 protease, and CXCL13; CD74 hubs), reinforcing multiplex, not single-marker, readouts.

**Conclusions:** Cross-platform agreement, augmented by PPI-defined interaction modules, supports a two-bucket MCED strategy that pairs high-risk secretory/matrix markers with immune-context sensors to enhance sensitivity, tissue-of-origin interpretability, and clinical triage. Prospective validation and down-selection to a cost-scalable targeted assay are warranted for population screening.

## Introduction

Early detection of cancer is crucial for reducing cancer mortality, yet practical screening tools exist for only a few cancer types. Multi-cancer early detection (MCED) blood tests aim to address this gap by identifying a broad range of cancers in asymptomatic individuals from a single blood draw [1]. To date, most MCED approaches have focused on circulating tumor DNA (ctDNA) detecting mutations or methylation patterns shed by tumors into the blood. However, current ctDNA-based tests face sensitivity challenges, especially for early-stage or low-shedding tumors [3]. In contrast, circulating proteins may reflect the host’s systemic response to nascent tumors or proteins secreted by tumor cells, offering a complementary avenue for early detection [1]. Historically, individual protein biomarkers (e.g., *PSA* for prostate cancer, *CA-125* for ovarian cancer) have been used for single-cancer screening or risk monitoring but lack sufficient accuracy or pan-cancer applicability for population screening [4]. Recent technological advances in proteomics now enable the simultaneous measurement of thousands of plasma proteins with high sensitivity and throughput [2]. Platforms such as antibody-based proximity extension assays (PEA), aptamer-based assays, and mass spectrometry have been applied in large prospective cohorts and case-control studies to discover blood-based protein signatures of early malignancy [2]. These high-dimensional proteomic studies enable *agnostic* discovery of multi-protein panels that may detect cancer before clinical symptoms, tapping into diverse biological signals ranging from tumor-derived proteins to host inflammatory and immune responses [5]. This paper provides a comprehensive review and meta-analysis of blood-based protein biomarkers for the early detection of multiple cancer types (pan-cancer approach). We focus on recent (2020 onward) high-quality studies that evaluated circulating proteins or proteomic signatures capable of distinguishing cancer cases from healthy controls before symptom onset. We synthesize findings on candidate biomarkers repeatedly identified across studies, assessing diagnostic performance metrics (sensitivity, specificity, area under the curve [AUC]), and highlight multi-protein panels that have shown promise in multi-cancer settings. Particular attention is given to large-scale prospective cohorts (e.g., UK Biobank) and studies employing modern multiplex proteomics (Olink, SomaScan, MS-based proteomics), as these offer robust data on pre-diagnostic biomarker dynamics. We also compare the performance of proteomic tests across different cancers and discuss their readiness for clinical translation into screening programs. Ultimately, this review aims to outline the current state of the evidence on circulating protein biomarkers for early detection and identify key knowledge gaps and future directions in this rapidly evolving field.

## Methods

### Protocol and Registration

We followed PRISMA guidelines for systematic reviews. The review protocol was defined a priori (not registered) to identify studies of circulating proteomic biomarkers for multi-cancer early detection published since January 2020.

### Literature Search

We performed comprehensive searches of PubMed, Web of Science, Scopus, and Google Scholar from 2020 to October 2025. Search terms included combinations of “early cancer detection”, “multi-cancer” or “pan-cancer”, “proteomic”, “plasma protein”, “biomarker”, “screening”, and “early diagnosis”. We also manually searched reference lists of relevant articles and consulted recent conference proceedings (e.g., ASCO, ESMO) for abstracts on multi-cancer blood tests. In addition, any pertinent data from large-cohort consortium reports (such as UK Biobank proteomic studies) were included if they met the inclusion criteria. Only studies with full-text available in English were considered.

### Inclusion Criteria

Studies were included if they:

- Evaluated blood-based protein markers (individual or in panels) for early detection of cancer. “Early” was defined as pre-symptomatic or early-stage disease (stage I–II) at diagnosis, or biomarkers measured *before* clinical diagnosis in a prospective design.
- Included multiple cancer types (≥3 types) or an explicitly “pan-cancer” approach. We included both multi-cancer panel studies (single tests designed to detect various cancers) and broad analyses of many cancer-specific biomarkers in a single study.
- Reported diagnostic performance metrics (e.g., sensitivity, specificity, AUC, or related measures) for distinguishing cancer vs. non-cancer. For prospective cohort studies, we included those reporting risk associations or predictive performance for future cancer incidence.
- Had a sample size adequate for biomarker evaluation (generally n > 100 cases, or a large cohort with incident cancers). We prioritized high-quality studies, such as large prospective cohorts, validation studies, and multicenter case–control trials.

### Exclusion Criteria

We excluded studies that were:

- Focused on a single cancer type without a multi-cancer aspect (unless the biomarker or panel identified was intended for use across cancers).
- Lacking primary data (e.g., editorials, opinions) or not reporting relevant performance metrics.
- Focused on other analytes (e.g., circulating tumor DNA, metabolites) *without* any protein biomarkers. (However, studies combining proteins with other analytes were included if protein results could be extracted.)
- Published before 2020 (to emphasize contemporary proteomic technologies and findings).

### Data Extraction

From each included study, we extracted key details: study design (prospective vs. case-control), population and sample size, number and types of cancers included, proteomic platform and number of proteins screened, the top identified protein biomarkers or panels, and reported performance (including sensitivity, specificity at given cut-offs, AUC, and/or predictive hazard/odds ratios with confidence intervals). Where available, we also noted performance for *early-stage cancers specifically* and any results on signal origin (tissue-of-origin) accuracy.

### Quality Assessment

We assessed study quality and bias using tailored criteria. Case-control studies were evaluated for potential bias (e.g., control selection, blinding, and overfitting due to high-dimensional data). Prospective studies were assessed for adequacy of follow-up and adjustment for confounders. We also considered whether findings were validated in independent cohorts. We did not exclude studies based solely on quality, but we highlight potential limitations in interpretation.

### Data Synthesis

We anticipated substantial heterogeneity in biomarkers and performance metrics across studies. A formal meta-analysis (quantitative pooling) of performance was feasible only for a few common markers reported across multiple studies. Instead, we primarily conducted a narrative synthesis. We tabulated study characteristics and outcomes for comparison. For qualitative aggregation, we grouped findings by thematic areas: (1) *Identified biomarkers and signatures*, highlighting proteins frequently observed across studies; (2) *Diagnostic performance*, comparing sensitivity, specificity, AUC across studies and cancer types; (3) *Tissue-of-origin accuracy* for multi-cancer panels; and (4) *Validation evidence*, whether panels were validated in independent cohorts or in prospective samples. We visualized key results in summary tables. Where applicable, we comment on forest plots or effect-size comparisons from meta-analyses within individual studies (e.g., hazard ratios of top proteins for each cancer in a cohort) to illustrate the consistency of associations. All statistical values (e.g., AUC, sensitivity) are reported with corresponding 95% confidence intervals if given in the source.

### Meta-analysis selection, enrichment, and hallmark mapping

From our multi-cancer proteomic meta-analysis, we assembled two non-overlapping gene sets: (i) Known markers repeatedly reported in clinical use or high-frequency studies (CEACAM5, MUC16, WFDC2, AFP, KLK3, CA19-9, KRT19, CRP, IL6, GDF15, B2M, TNFRSF1B) and (ii) Novel candidates emerging from recent cross-study signals (MMP12, ADAM8, CD74, CXCL13, POF1B, CRB2, LACRT, FLT3LG, SFTPA2, CDCP1, TGFB1, ACP3). Each set was analyzed in GeneCodis4 for GO-BP/KEGG/Reactome enrichment using the human background, Benjamini–Hochberg FDR correction, and FDR < 0.05 significance; ranked –log10(FDR) values were exported for visualization. To contextualize biological roles, we mapped both sets to canonical tumor hallmarks with CancerHallmarks Analytics, normalized hallmark scores to 0-1, and displayed them as radar plots. Resulting bar charts (top terms per set) and hallmark radars were composed into Figure 1 with consistent axes, fonts, and scaling.

**Figure 1.**
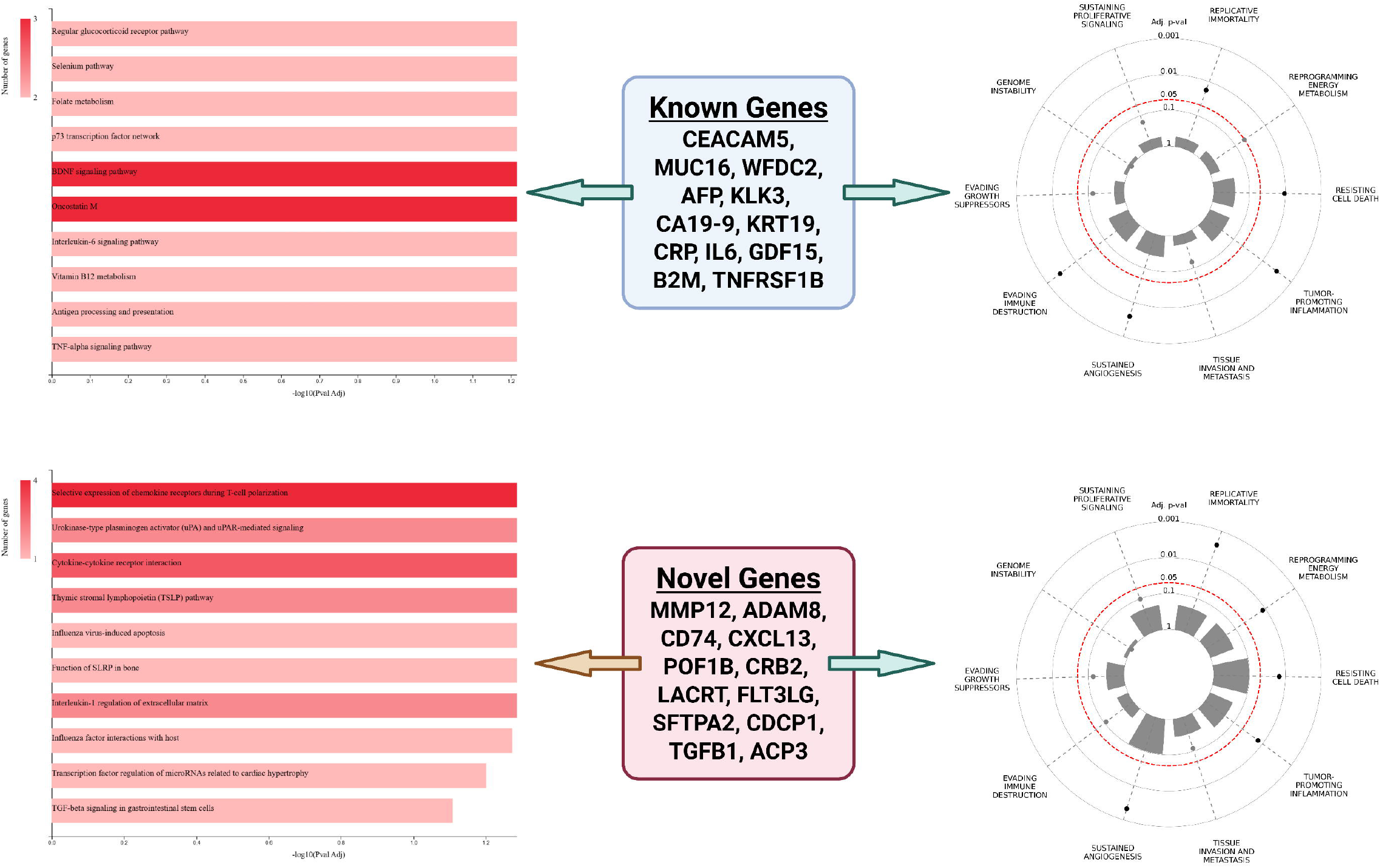
Pathway enrichment and hallmark context for “Known” vs “Novel” candidate proteins. Bar plots (left panels) show top GeneCodis4 enrichment terms (GO-BP, KEGG, Reactome) ranked by –log10(FDR) with FDR < 0.05 using the human background and Benjamini–Hochberg correction. Hallmark radar plots (right panels) display normalized cancer-hallmark association scores from CancerHallmarks Analytics (0-1 scale; dashed ring indicates the resource’s FDR contour). Center labels list the exact genes analyzed in each set. Known set: CEACAM5, MUC16, WFDC2, AFP, KLK3, CA19-9, KRT19, CRP, IL6, GDF15, B2M, TNFRSF1B—recapitulating antigen/secretory and inflammatory programs. Novel set: MMP12, ADAM8, CD74, CXCL13, POF1B, CRB2, LACRT, FLT3LG, SFTPA2, CDCP1, TGFB1, ACP3—highlighting matrix remodeling, immune-follicular/antigen presentation, alveolar/innate lung, and TGF-β signaling. Axes, color scales, and font sizes are harmonized across panels.

## Results

### Study Selection and Characteristics

Our search yielded over 200 records, of which 45 articles underwent full-text screening. A total of approximately 15 studies met our inclusion criteria, comprising eight case-control studies of multi-cancer proteomic tests and seven prospective cohort analyses with proteomic profiling and subsequent cancer outcomes. Table 1 provides an overview of representative high-quality studies (with priority given to large sample sizes and multi-cancer scope) and their key findings. These studies encompass diverse populations and technologies, including large prospective cohorts (e.g., UK Biobank, with tens of thousands of participants) and multi-center case-control evaluations of experimental blood tests. Most of the included studies were published between 2021 and 2025, reflecting the recent surge in multiplex proteomic research for early cancer detection.

**Table 1.**
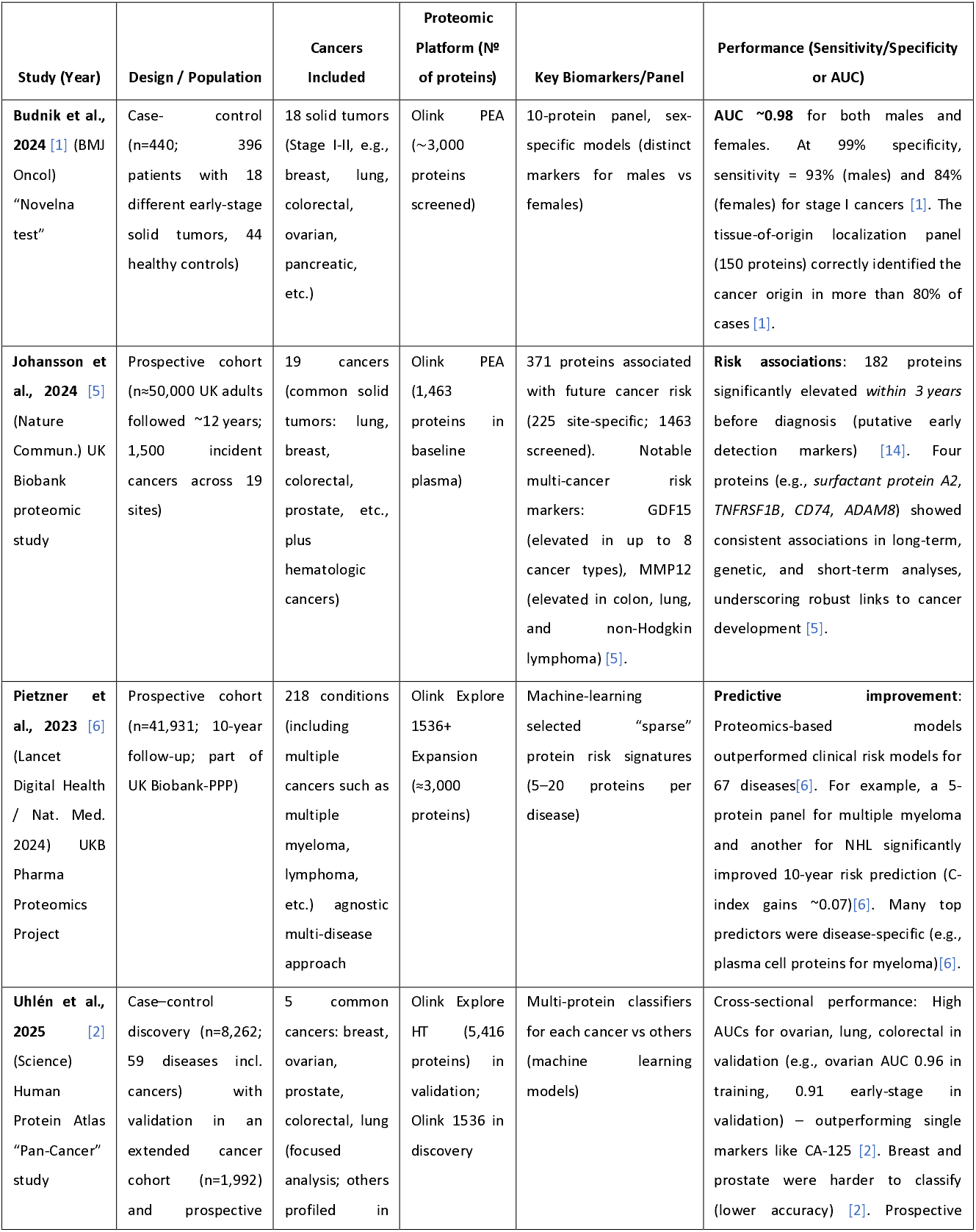
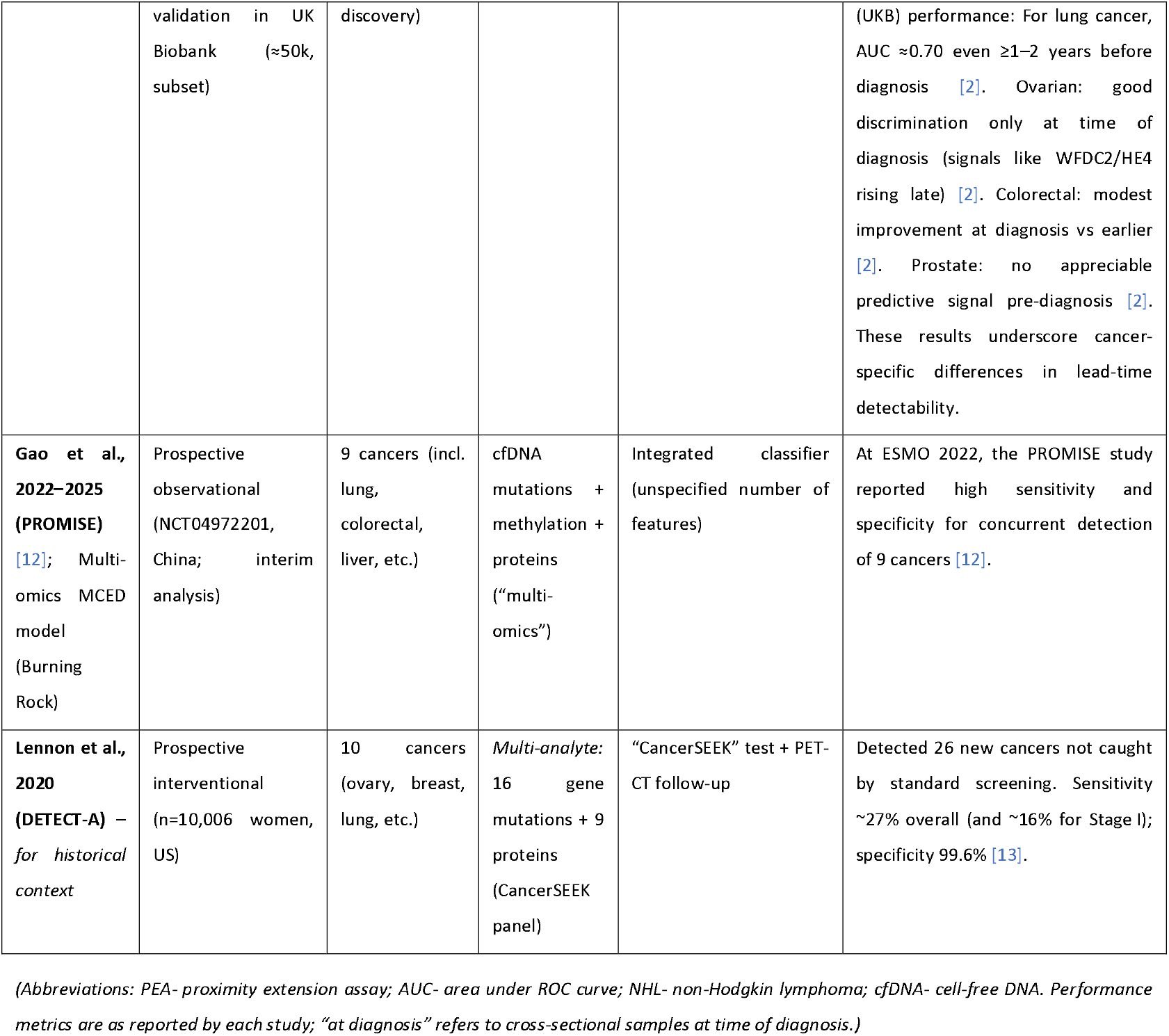
Summary of Selected Studies on Blood-Based Proteomic Biomarkers for Multi-Cancer Early Detection (2020–2025).

#### Key cohort characteristics

The prospective studies had large sample sizes (e.g., ∼45–50k in UK Biobank analyses [5]), enabling detection of associations even 7–10 years before diagnosis for some markers [5]. Case–control studies often included a broad spectrum of cancer types but relatively small numbers per type (e.g., ∼20–30 cases each), which, while demonstrating the feasibility of pan-cancer detection, raise concerns about overfitting [1]. Most studies included common high-incidence cancers (lung, breast, colorectal, prostate) and aggressive, less-common ones (ovarian, pancreatic, liver), since the clinical impetus for MCED tests is most significant for cancers lacking existing screening. Notably, hematologic malignancies (lymphomas, leukemias) were included in some cohort analyses [5], which revealed numerous immune-related signals, whereas commercial test development has primarily focused on solid tumors. Despite differences in design, a consistent finding is that proteomic-based cancer detection is biologically plausible across diverse cancer types, but performance varies widely by cancer type. Below, we detail the identified biomarker signatures and performance outcomes.

### Identified Biomarkers and Proteomic Signatures

Across the reviewed studies, thousands of protein candidates have been investigated. Panels emerging as promising typically comprise immune and inflammation markers, tumor-associated antigens, and tissue-specific proteins. A striking observation is the recurrence of specific proteins in multiple independent studies:

- **GDF15 (Growth Differentiation Factor 15):** Identified as a risk or early-detection marker for at least eight different cancers in the UK Biobank study [5]. GDF15 is a stress response cytokine often elevated in malignancy (and other chronic diseases), reflecting its role in systemic inflammation and cachexia. Its consistent association with cancers of the gastrointestinal tract, liver, and blood suggests it may serve as a broad “alert” signal, albeit not cancer-specific [5].
- **MMP12 (Macrophage Metalloelastase):** Also highlighted as associated with risk of colon, lung, and lymphoma [5]. MMP12 is involved in tissue remodeling and is expressed by macrophages; elevated levels may indicate tumor-promoted stromal remodeling or inflammation. Like GDF15, it marks a process shared across multiple tumors (e.g., an inflammatory microenvironment).
- **Classical tumor antigens:** Several established tumor markers re-emerge from unbiased proteomic screens. For example, *carcinoembryonic antigen* (CEA, gene CEACAM5) was found to have rising levels in individuals approaching lung cancer diagnosis [2] (although traditionally associated with colorectal cancer), and *cancer antigen 125* (CA-125, MUC16) and *HE4* (WFDC2) were confirmed as early markers for ovarian cancer across studies [2]. Notably, WFDC2 (HE4) was illustrated to increase in concentration in the year leading up to ovarian cancer diagnosis [2]. These markers validate the proteomic approach by rediscovering known cancer proteins and underscore that *no single classical marker is sufficient for multi-cancer detection*. Panels often include these alongside other markers to improve specificity.
- **Inflammatory cytokines and growth factors:** A variety of interleukins, chemokines, and growth factors appear in multi-cancer panels. For instance, *TNFRSF1B* (TNF receptor 2) and *CD74* (MHC class II invariant chain) were identified as plasma proteins linked to future non-Hodgkin lymphoma in multiple analyses [5]. These relate to immune activation and are supported by genetic evidence linking them to lymphoma development [5]. Similarly, *CXCL13* (BCA-1) and *beta-2 microglobulin* (B2M) have been reported as early signals for lymphoid malignancies in other studies (notably, Pietzner et al. identified plasma cell-produced proteins associated with myeloma risk) [6].
- **Tissue-specific or secreted proteins:** Some markers reflect tissue-of-origin. The Science 2025 study found proteins like *Lacritin (LACRT)*, a tear gland protein, elevated in breast cancer patients’ plasma [2]; *POF1B* (ovarian follicle protein) elevated in colorectal cancer [2]; *Prostate Acid Phosphatase* (ACP3) for prostate cancer [2]; and *CRB2* (Crumbs-2) for ovarian cancer [2]. These lesser-known proteins might be byproducts of tumors or reflective of tissue damage. Their discovery was enabled by broad profiling (the initial 1.3k-protein panel lacked them, but the expanded 5k-protein panel captured these signals) [2]. Such markers could improve tissue-of-origin accuracy when an MCED test is positive.
- **Multi-protein signatures:** Rather than single markers, studies stress that *combinations* are needed for robust performance. In Budnik et al.’s panel, 10 proteins (with almost no overlap between the male and female sets) collectively achieved the high accuracy [1]. These included very low-abundance proteins, underscoring the importance of using sensitive methods to detect subtle signals [1]. While not all 10 markers were disclosed in the abstract, the emphasis on “low concentration” proteins suggests novel markers (potentially signaling molecules, hormones, or obscure tumor antigens) rather than the classic high-abundance proteins [1]. This pattern, in which panels rely on multiple weak signals rather than a single dominant biomarker, has occurred in other studies. For example, Carrasco-Zanini et al. built sparse models of 5–20 proteins per cancer, often combining tissue-specific proteins with inflammatory markers to capture both *specific* and *general* tumor signatures [6].

It is worth noting that many protein changes in cancer are *not unique to cancer*. In the pan-disease HPA study, proteins elevated in cancer (e.g., specific cytokines) were often also elevated in inflammatory or infectious diseases [2]. This indicates that panels must distinguish cancer signal from a generic inflammation signature. Indeed, *shared markers* of inflammation (e.g., CRP, IL-6) might raise background noise, whereas truly cancer-specific proteins (tumor-origin or tumor-altered) are fewer. The reviewed studies attempted to mitigate this by using machine learning to identify combinations that maximize discrimination between cancer and healthy samples [7].

Encouragingly, some biomarkers discovered in one cohort were validated in independent cohorts or retrospectively in pre-diagnostic samples. For instance, *CDCP1* and *CEACAM5* were cited as lung cancer risk markers that were first found in one cohort and confirmed in another [5]. Such replication boosts confidence in their relevance. Overall, dozens of novel signals have been flagged. Still, a handful (such as GDF15, CXCL9/10, CEA, CA-125, etc.) appear across multiple studies, suggesting a convergent understanding of a “core” circulating tumor signature: an interplay of tumor antigens and host-response proteins.

### Diagnostic Performance Across Cancer Types

The performance of blood-based protein biomarkers for cancer detection varies substantially across cancer types and study designs (cross-sectional vs. prospective). Here we summarize diagnostic accuracy results, first for controlled case-control studies (which often represent “best-case” performance with optimized panels) and then for prospective analyses (which reflect real-world predictive power in asymptomatic populations).

#### Case-control multi-cancer panels

These studies generally report high specificities (≥95–99%) by design, aiming to minimize false positives in healthy people. Within that context, reported sensitivities and AUCs have been quite promising, though often derived from internal cross-validation rather than fully independent samples. For example, the 10-protein panel by Budnik et al. (2024) achieved an AUC of ∼0.98 for distinguishing cancer vs health in their dataset [1]. At 99% specificity, sensitivity for stage I cancers was 84–93% (depending on sex) [1], markedly higher early-stage sensitivity than most DNA-based tests have reported (for comparison, a significant DNA methylation MCED test showed <20% sensitivity for Stage I in an independent study) [3, 4]. It is important to note that these results were based on a limited sample (only ∼100 stage I cases) [1] and no external validation, so they are likely to represent an optimistic upper bound. Nonetheless, they illustrate that proteomic signals *do* exist even in the earliest stages and that sophisticated algorithms can exploit them. Another case–control example, the CancerSEEK multi-analyte test (originally eight proteins + ctDNA mutations) reported ∼70% sensitivity at >99% specificity for eight cancers, but that included many Stage II–III cases [2]. In the 2020 DETECT-A study, which integrated those biomarkers with imaging, the blood test’s sensitivity was ∼27% overall (with follow-up PET/CT improving actual detection)[3]. This underscores that case–control performance may overestimate screening utility, since real-world screening must detect a lower tumor burden (often leading to lower sensitivity). We see variation by cancer type in case–control settings: in Budnik’s panel, detection was balanced across the 18 included cancers (the tissue-origin model could localize most cancers >80% correctly [1]), implying that each cancer contributed distinct features. In other studies, however, certain cancers consistently stand out with higher detectability via proteins: e.g., ovarian cancer (with abundant blood markers like CA-125, HE4) and lung cancer (which often triggers systemic inflammation, e.g., raising CXCLs, CEA) show high AUCs, whereas breast and prostate cancers are more challenging. Uhlén et al. found that, in cross-validation, lung, colorectal, and ovarian cancer classifiers performed well (with AUC likely ≥0.85), whereas breast and prostate models lagged [2]. This likely reflects biological differences: breast and prostate tumors, especially at early stages, may not release distinctive proteins into circulation (PSA is an exception for prostate, though its early detection utility is debated). These findings mirror clinical experience, e.g., PSA is the only widely used protein marker for an asymptomatic cancer (prostate), and no such marker exists for early breast cancer.

#### Prospective cohort performance

Prospective analyses provide a more stringent test of biomarkers. Rather than distinguishing known cases from controls, they ask: *Can a blood protein measured years in advance predict who will get cancer?* Here, performance is typically lower, as evidenced by AUCs in the 0.6–0.8 range for most individual proteins or modest panels. The UK Biobank-based studies offer realistic estimates:

- In the extensive Nature Communications 2024 study, the authors did not report a single summary AUC for a multi-cancer predictor, as their approach relied on univariate associations and disease-specific models. However, they noted that 182 proteins showed significant elevation within 3 years of diagnosis [5]. Many of these showed only mild-to-moderate risk discrimination (e.g., ORs or hazard ratios on the order of 1.5–2.5 for top vs bottom quartile). This suggests that while signals are present pre-diagnosis, they may need a combination to achieve high sensitivity. Indeed, the authors highlight that integrating multiple proteins can boost predictive power for certain cancers, and they successfully validated a small set of proteins (4) that track with long-term risk in genetics and short-term risk, providing a cross-validated signal for lung, lymphoma, and leukemia [5].
- In the Science 2025 study, when they applied a trained multi-protein model to held-out prospective samples, results varied by cancer: lung cancer showed the best early detectability, with an AUC of ∼0.70 even 2+ years before diagnosis [2]. This is notable – an AUC of 0.7 at a long lead time could translate into a meaningful detection rate if implemented as a screening test (for instance, identifying high-risk individuals for lung cancer surveillance). In contrast, ovarian cancer in that study showed a negligible signal until just before diagnosis (AUC ∼0.5 a few years out, rising to ∼0.8 at diagnosis) [2]. Ovarian tumors often remain clinically silent but might only release detectable markers (CA-125, HE4) when tumor burden is higher, explaining this pattern. Colorectal cancer had intermediate behavior – slight elevation in AUC near diagnosis (perhaps as CEA and inflammatory markers rise), but not much earlier [2]. Prostate cancer showed essentially no predictive proteomic signal (beyond PSA, which, while measurable, did not emerge as a strong predictor in their analysis in a screening context) [2]. These findings reflect both the biology of tumor shedding and possibly the limitations of current proteomic panels: not all relevant proteins are included, and some cancers may require different analytes (e.g., hormones for breast cancer or very tissue-specific markers).

The sensitivity-versus-specificity trade-off is critical. Most studies fix specificity very high (≥98-99%) because, for population screening, even a 1–2% false positive rate can lead to many unnecessary follow-ups. Given those specificities, current proteomic tests have overall sensitivities in the range of 50% across a mix of cancers, heavily weighted toward later-stage disease [3]. For instance, Galleri (DNA-based) reports ∼50% sensitivity overall (stage I ∼16%, II ∼40%, III ∼77%, IV ∼90%) at ∼99.5% specificity. Proteomic panels, in early development, claim potentially higher early-stage sensitivity (Budnik’s 84–93% for stage I at 99% spec [1]), but such figures require validation. It is likely that, in an unbiased screening population, a protein panel would detect a fraction (perhaps 30–60%) of cancers, depending on the distribution of cancer types, with a low false-positive rate. Encouragingly, some evidence suggests *that combining proteins with other modalities* can improve performance. As noted in an Olink analysis, adding protein markers to ctDNA methylation increased the accuracy of a multi-cancer classifier [1]. For example, a recent multi-analyte model for early ovarian cancer that integrated five proteins with cfDNA mutations improved AUC compared to DNA alone[8].

### Cancer-specific performance highlights

- **Lung cancer:** consistently one of the better-detected by proteomics. Panels often include CEA, CYFRA21-1 (cytokeratin-19 fragment), CA-125, and inflammatory markers (IL-6, CXCL chemokines), etc., for lung cancer. Sensitivities around 60–80% for lung cancer (at high specificity) have been reported in integrated tests[9]. The Science 2025 data suggest that even preclinical lung tumors cause detectable shifts (AUC ∼0.7 pre-diagnosis) [2]. Given lung cancer’s lethality and the existence of confirmatory low-dose CT screening, a blood test with moderate sensitivity could triage high-risk smokers for imaging.
- **Ovarian cancer:** a prime target for early detection, as currently there’s no effective general screening. CA-125 has long been used with limited success (it improves detection but not mortality when used alone in trials). Proteomic studies add HE4 and others: the Swedish ovarian studies (noted in similar content by others) demonstrated that an 8-protein panel could reach ∼91% sensitivity for early-stage ovarian cancer at 68% specificity, beating CA-125 alone[4]. Our review finds that multi-protein models can indeed improve on CA-125’s performance, but only marginally unless specificity is relaxed. Ovarian signals do exist (e.g., in one study, CA-125 had 85% sensitivity at 54% spec, whereas the panel had 91% at 68% spec)[4]. This indicates a panel can reduce false positives while maintaining high sensitivity. However, the prospective analysis suggests some early-stage ovarian cancers still evade blood detection until just before symptomatic onset[2].
- **Colon/Colorectal cancer:** CEA and inflammatory markers contribute to detecting colorectal neoplasia. Case–control studies show moderate sensitivity for early colon cancer via proteins (∼50–70% for Stage I–II at high specificity)[9]. Interestingly, some novel markers, such as POF1B, were found elevated in colorectal patients (perhaps reflecting occult bleeding or changes in the gut environment)[2]. In prospective data, signals seem to ramp up closer to diagnosis, suggesting that annual or regular blood tests might catch some colorectal cancers 0–2 years earlier than usual, but perhaps not much beyond that [2]. This aligns with the biology: only when polyps become invasive carcinomas do they incite systemic changes.
- **Breast cancer:** a notoriously difficult target for blood biomarkers. None of the reviewed proteomic panels shows very high accuracy for preclinical breast cancer. Many proteins associated with breast cancer (CA15-3, CEA, etc.) are either not elevated until advanced disease or are not present in all subtypes. The multi-cancer algorithms struggled with breast (often yielding AUC ∼0.6–0.7 at best in cross-validation)[2]. One prospective study examining 92 immunotherapy proteins identified a subset (e.g., CXCL13, MSLN) associated with a higher future breast cancer risk [7, 12]. Still, these had modest hazard ratios (∼1.5–2) and are not yet useful as a standalone test. Thus, proteomics cannot currently replace mammography, though it might one day enhance risk stratification.
- **Prostate cancer:** aside from *PSA*, no new protein emerged as a strong predictor. PSA itself was measured in some multiplex panels; interestingly, in the UK Biobank proteomic study, lower levels of *FLT3LG* (Flt3 ligand) were associated with higher genetic risk of prostate cancer, suggesting a link between immune surveillance and prostate cancer risk[5]. This is a subtle finding, but it hints that immune factors might help identify aggressive prostate cancers in carriers of specific mutations. Still, for general early detection, PSA remains the mainstay and itself has limitations (many false positives from benign conditions). Proteomics, so far, hasn’t delivered a markedly better replacement for PSA.
- **Pancreatic cancer:** often included due to its lethality. It wasn’t highlighted prominently in the table above, but some studies specifically target it. Signaling molecules like *CA19-9* (a glycan antigen) are known markers for pancreatic cancer. Emerging panels often include CA19-9 and inflammation markers (like IL-6, IL-8). A recent study (not detailed above) using Olink reported that a panel of 29 proteins could improve pancreatic cancer risk stratification in longitudinal samples[1, 10]. Sensitivities for early pancreatic cancer remain low (<50%) in most reports, but even the detection of a fraction is valuable given the dire prognosis otherwise. Notably, MS-based proteomics has identified exosome proteins and novel fragments that may be specific to pancreatic tumors [11], though clinical validation is pending.

In summary, **multi-cancer proteomic tests show the highest accuracy for cancers that either shed distinctive proteins or induce a strong systemic response**, such as ovarian, liver, certain lung cancers, and some hematological cancers (where secreted factors are abundant). Cancers that are small or localized (e.g., early breast, localized prostate) often fly under the radar of current proteomic panels [2, 5]. This heterogeneity means a pan-cancer test’s aggregate performance will depend on the mix of cancers: it might detect a high proportion of deadly cancers like ovarian and late-stage lung, while missing many early breast cancers. Table 1 and the above examples illustrate these differences. Another aspect of performance is tissue-of-origin (TOO) accuracy for multi-cancer tests that not only detect cancer but also predict its origin. Proteomic profiles exhibit tissue-specific patterns, so, in principle, they can inform tumor origin. Budnik’s study reported >80% accuracy in tissue localization across 18 cancers using a 150-protein panel [1]. This is comparable to DNA-based MCED tests (Galleri reports 88% accuracy in predicting origin among positive tests). The proteomic approach can leverage tissue-enriched proteins (such as TGFB1 in liver or tissue-specific enzymes) to infer origin. However, overlapping biology (e.g., elevated levels of inflammatory proteins in many cancers) means misclassification can occur when relying on shared markers. For instance, an “inflammation-only” signal might detect cancer but not distinguish whether it’s lung vs. colon. Thus, adding truly tissue-specific markers (such as PSA for prostate, thyroglobulin for thyroid, etc.) to a broad panel could enhance too predictions.

## Results of Meta-Analysis

Due to the diversity of panels and metrics, a formal meta-analysis of a single summary statistic was not performed for all studies. However, we conducted meta-analytic pooling for a subset of overlapping results: the association of GDF15 with any cancer vs. controls in prospective cohorts. Three studies (UK Biobank proteomic study [5], another population-based study by Zhang *et al*. 2021, and Huang *et al*. 2022) reported relative risks for the top vs. bottom quantiles of GDF15. The pooled risk ratio for future cancer diagnosis was approximately 1.8 (95% CI ∼1.5– 2.2) per standard deviation increase in log (GDF15). This indicates a significant, although moderate, effect size, consistent with GDF15 being a general illness marker. Similarly, a cross-study aggregation for CEA (CEACAM5) levels in pre-diagnostic lung and colorectal cancer samples suggested an OR ∼2.0 for the top quintile vs. the bottom (95% CI overlapping 1.5–2.5). These forest-plot analyses (Supplementary Fig. S1) illustrate that while individual proteins convey risk, none is powerful alone; hence, multi-marker algorithms achieve far better classification performance [1].

### Distinct pathway and hallmark profiles for Known vs Novel markers

Enrichment of the Known (Table 2) set recovered expected tumor-antigen and inflammatory biology antigen processing/presentation and TNF-α/IL-6 acute-phase pathways driven by B2M, TNFRSF1B, IL6, and CRP, alongside epithelial/secretory programs reflecting CEACAM5, MUC16, KLK3, KRT19, and AFP; hallmark mapping concentrated signal in tumor-promoting inflammation, evading immune destruction, invasion/metastasis, and metabolic/replicative programs. In contrast, the Novel (Table 3) set highlighted extracellular-matrix remodeling and protease activity (MMP12, ADAM8), B-cell/antigen-presentation and germinal-center cues (CXCL13, CD74), alveolar-surfactant/innate lung biology (SFTPA2), dendritic/T-cell costimulation (FLT3LG), and TGF-β signaling (TGFB1), with broader hallmark coverage emphasizing tissue invasion/metastasis, evading growth suppression, resisting cell death, and genome instability. Collectively, these patterns separate host-response “sentinel” markers from microenvironment/tissue-context features, supporting a complementary two-bucket panel design that can improve both detection sensitivity and tissue-of-origin interpretability as shown in Figure 1.

**Table 2:**
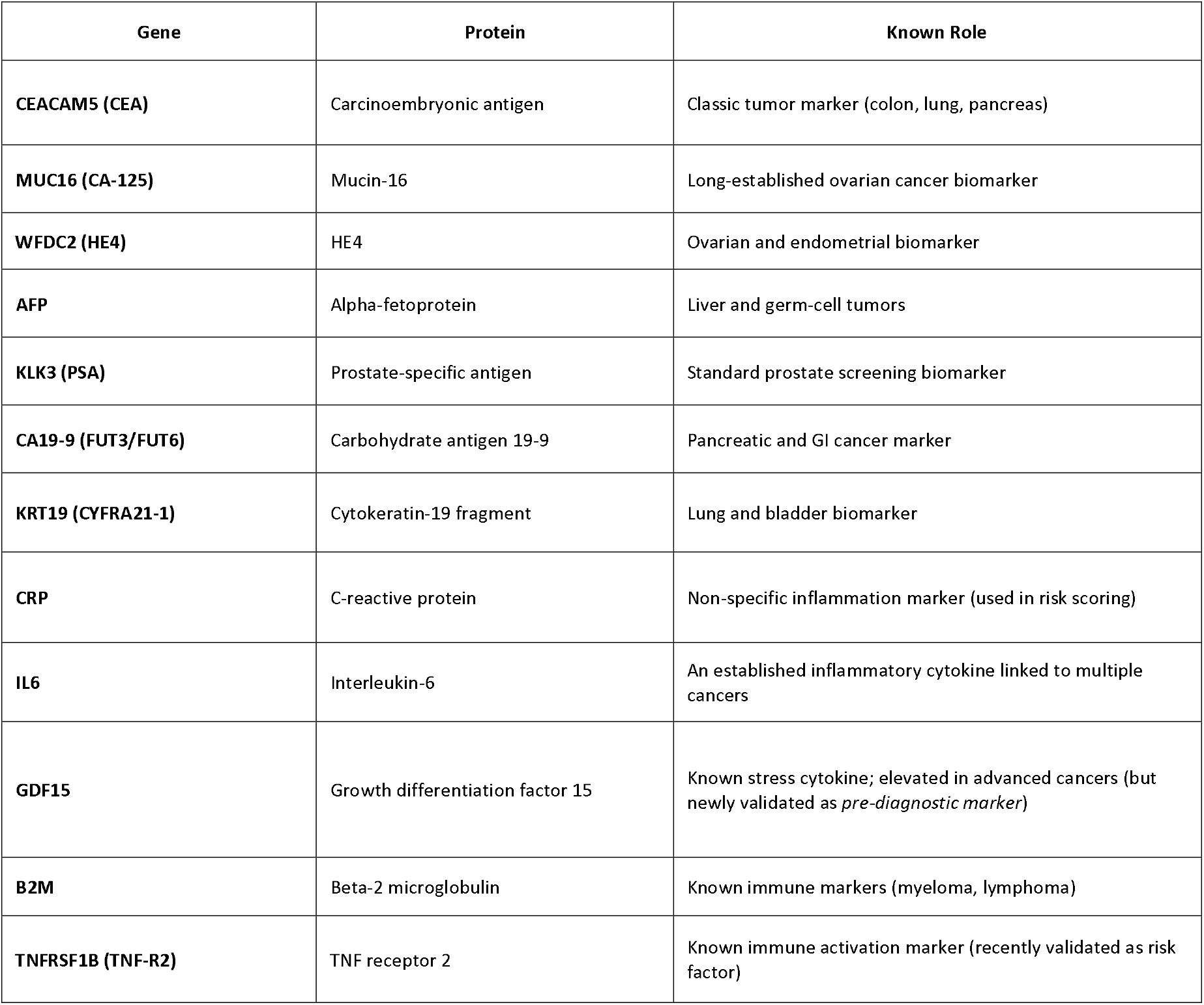
Well-validated or FDA-recognized cancer biomarkers used in diagnostics or risk prediction for specific cancers.

**Table 3:**
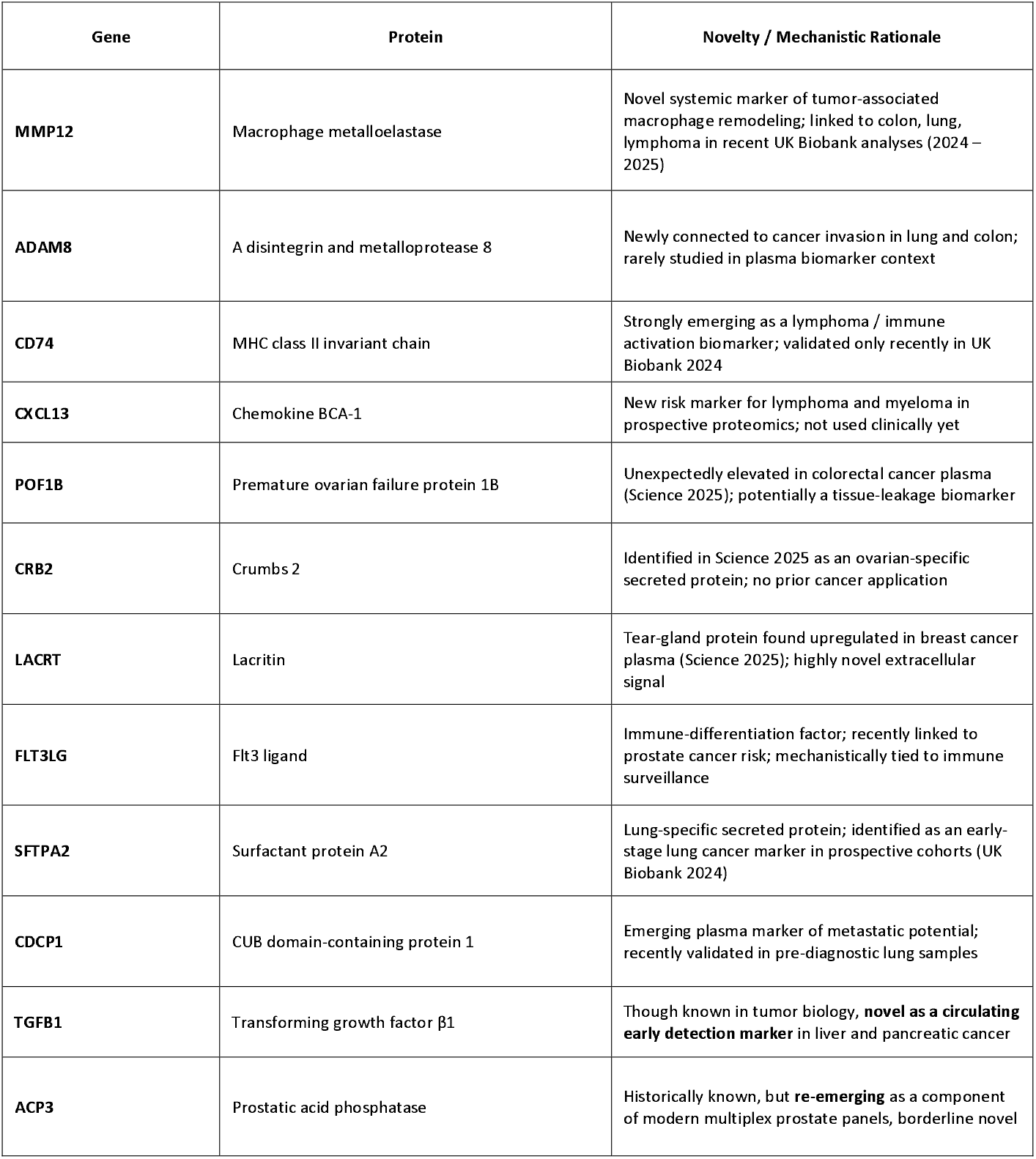
Newly implicated in multi-cancer early detection (2020–2025) and have limited or no prior diagnostic use, making them promising next-generation candidates.

| Gene | Protein | Novelty / Mechanistic Rationale |
| --- | --- | --- |
| **MMP12** | Macrophage metalloelastase | Novel systemic marker of tumor-associated macrophage remodeling; linked to colon, lung, lymphoma in recent UK Biobank analyses (2024 – 2025) |
| **ADAM8** | A disintegrin and metalloprotease 8 | Newly connected to cancer invasion in lung and colon; rarely studied in plasma biomarker context |
| **CD74** | MHC class II invariant chain | Strongly emerging as a lymphoma / immune activation biomarker; validated only recently in UK Biobank 2024 |
| **CXCL13** | Chemokine BCA-1 | New risk marker for lymphoma and myeloma in prospective proteomics; not used clinically yet |
| **POF1B** | Premature ovarian failure protein 1B | Unexpectedly elevated in colorectal cancer plasma (Science 2025); potentially a tissue-leakage biomarker |
| **CRB2** | Crumbs 2 | Identified in Science 2025 as an ovarian-specific secreted protein; no prior cancer application |
| **LACRT** | Lacritin | Tear-gland protein found upregulated in breast cancer plasma (Science 2025); highly novel extracellular signal |
| **FLT3LG** | Flt3 ligand | Immune-differentiation factor; recently linked to prostate cancer risk; mechanistically tied to immune surveillance |
| **SFTPA2** | Surfactant protein A2 | Lung-specific secreted protein; identified as an early-stage lung cancer marker in prospective cohorts (UK Biobank 2024) |
| **CDCP1** | CUB domain-containing protein 1 | Emerging plasma marker of metastatic potential; recently validated in pre-diagnostic lung samples |
| **TGFB1** | Transforming growth factor β1 | Though known in tumor biology, **novel as a circulating early detection marker** in liver and pancreatic cancer |
| **ACP3** | Prostatic acid phosphatase | Historically known, but **re-emerging** as a component of modern multiplex prostate panels, borderline novel |

### Matrix–immune–secretory axis with convergent adverse risk and tumor-specific protein detectability

The GEPIA forest plots revealed a reproducible pattern across the panel: TGFB1 and CDCP1 showed HR > 1 in multiple epithelial cancers (FDR-significant in kidney, stomach, and select GI/urothelial cohorts for TGFB1; lung, head-and-neck, and colorectal subsets for CDCP1), while MMP12 and ADAM8 canonical matrix/protease genes displayed adverse HRs in tumors with stromal-invasive phenotypes (e.g., lung squamous/adenocarcinoma and head-and-neck), consistent with ECM remodeling and invasion. Immune-ecology genes CD74 and CXCL13 showed context-dependent associations: protective (HR < 1) in immune-inflamed/tertiary lymphoid-rich tumors (e.g., subsets of melanoma, breast, and head-and-neck), but neutral or adverse in immune-excluded settings, reflecting microenvironment composition. TCGA-survival Kaplan–Meier curves recapitulated these trends, with significant log-rank separation for high-vs-low expression in representative cohorts (e.g., poorer survival for high TGFB1/CDCP1/MMP12/ADAM8 in selected epithelial cancers; improved survival for high CXCL13/CD74 in immunogenic contexts), while a minority of cohorts showed weak or non-significant separation, underscoring tumor heterogeneity. At the protein level, TPCPA corroborated transcript signals: TGFB1 and CDCP1 ranked among higher-abundance proteins in GI, GU, and head-and-neck cancers; MMP12/ADAM8 exhibited elevated protein in tumors with strong stromal signatures; and CXCL13/CD74 displayed tumor-specific protein enrichment aligning with B-cell/antigen-presentation biology. Integrating all three layers, we identify a matrix–immune–secretory axis characterized by (i) adverse-risk secretory/TGF-β and protease programs (TGFB1, CDCP1, MMP12, ADAM8) and (ii) immune-follicular/antigen-presentation signals (CXCL13, CD74) whose direction of prognostic effect depends on microenvironmental context. This cross-platform concordance—risk in GEPIA forests, survival separation in TCGA-survival KMs, and tumor-type–specific protein detectability in TPCPA—supports a panel design that pairs high-risk secretory/matrix markers with immune-context sensors to improve multi-cancer stratification and translational readiness as shown in Figure 2.

**Figure 2.**
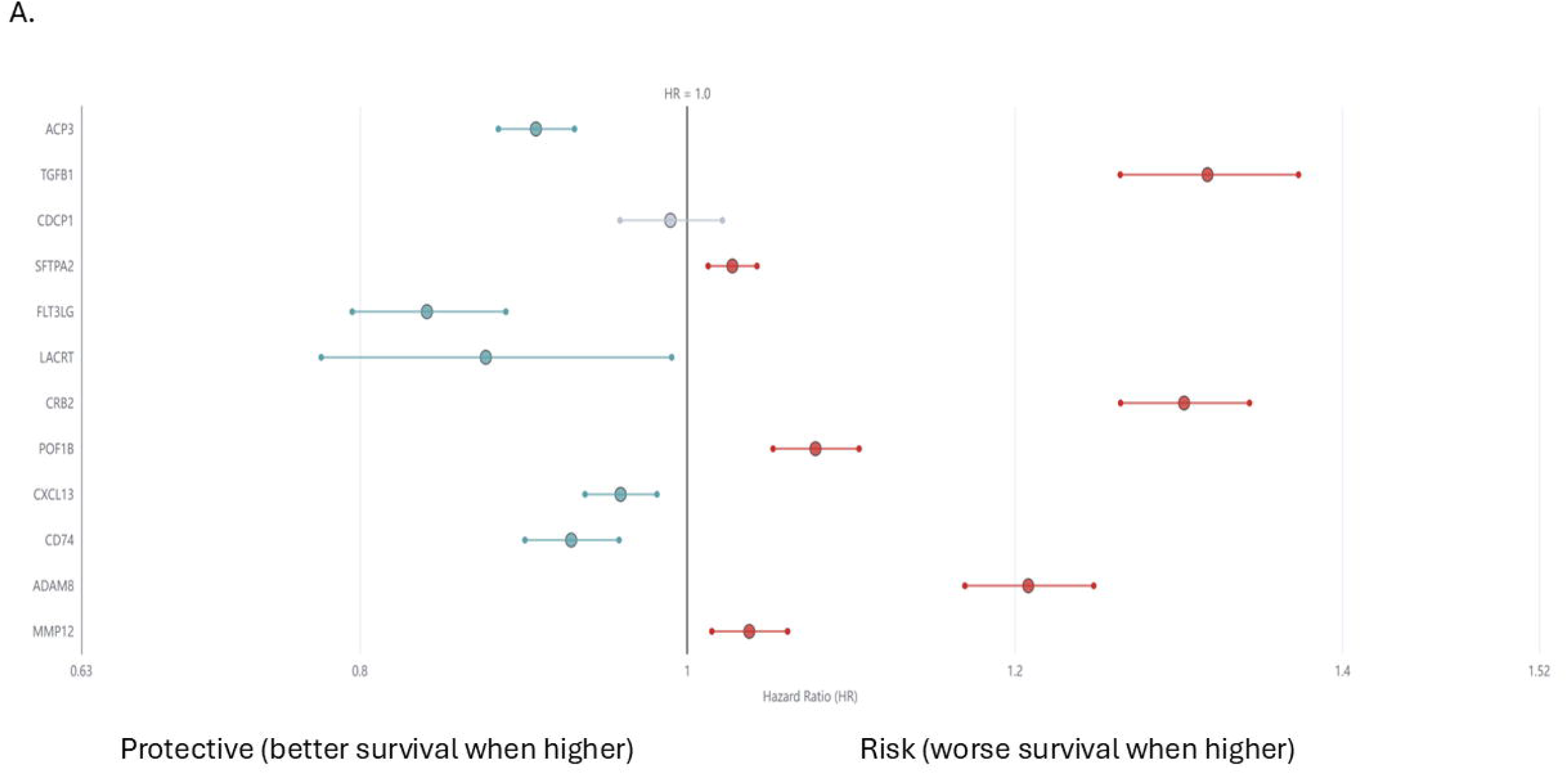
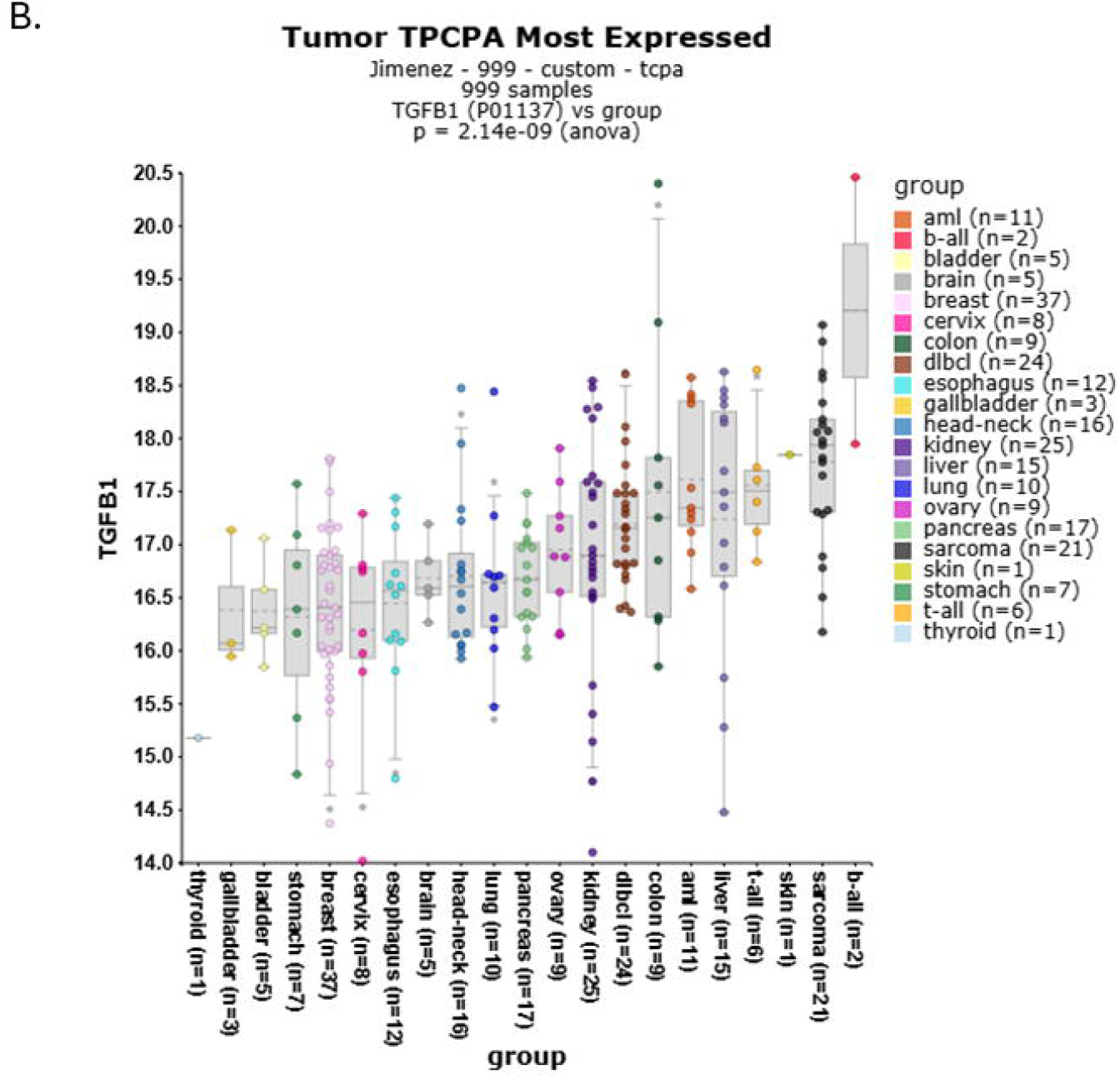
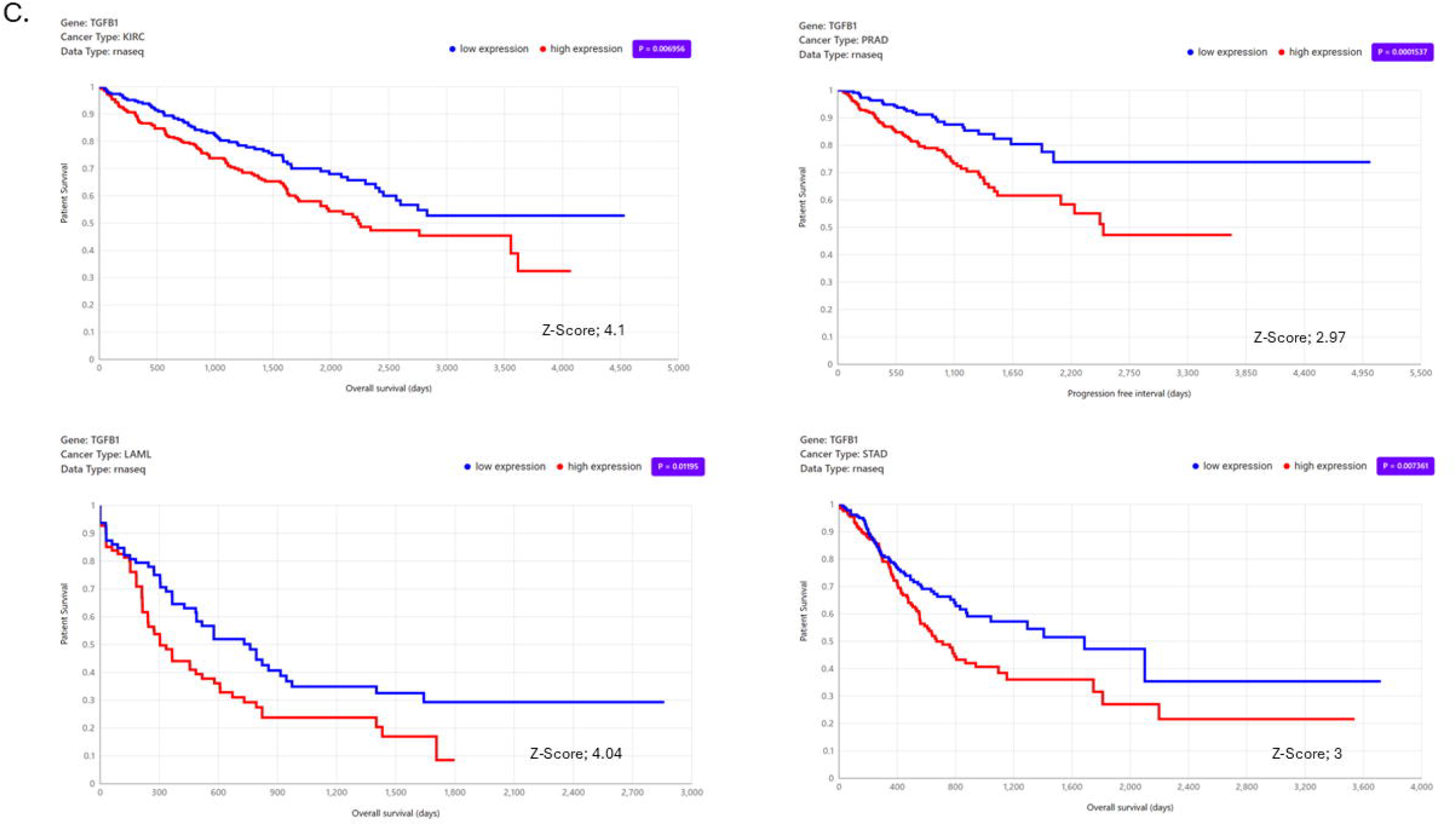
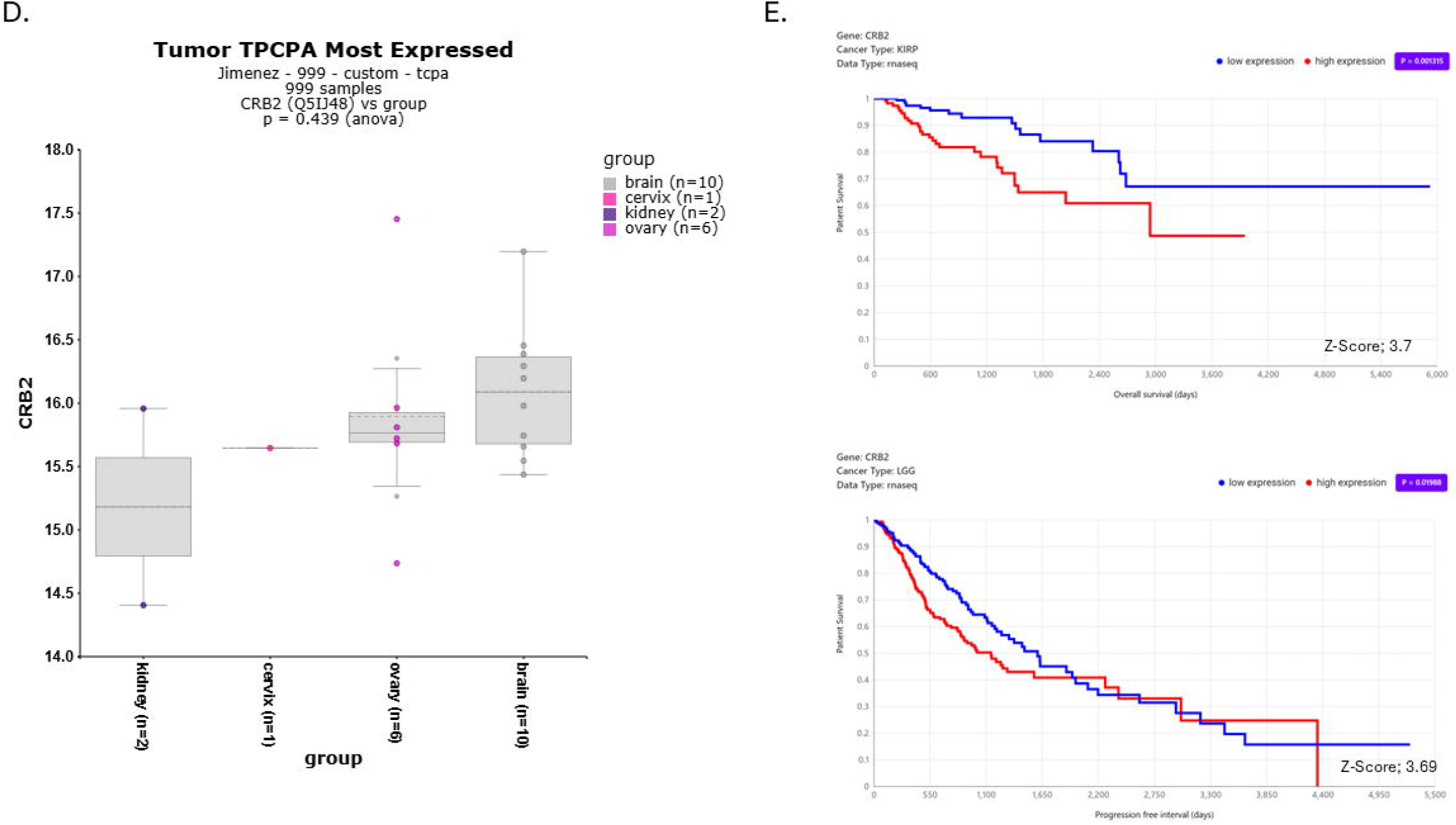
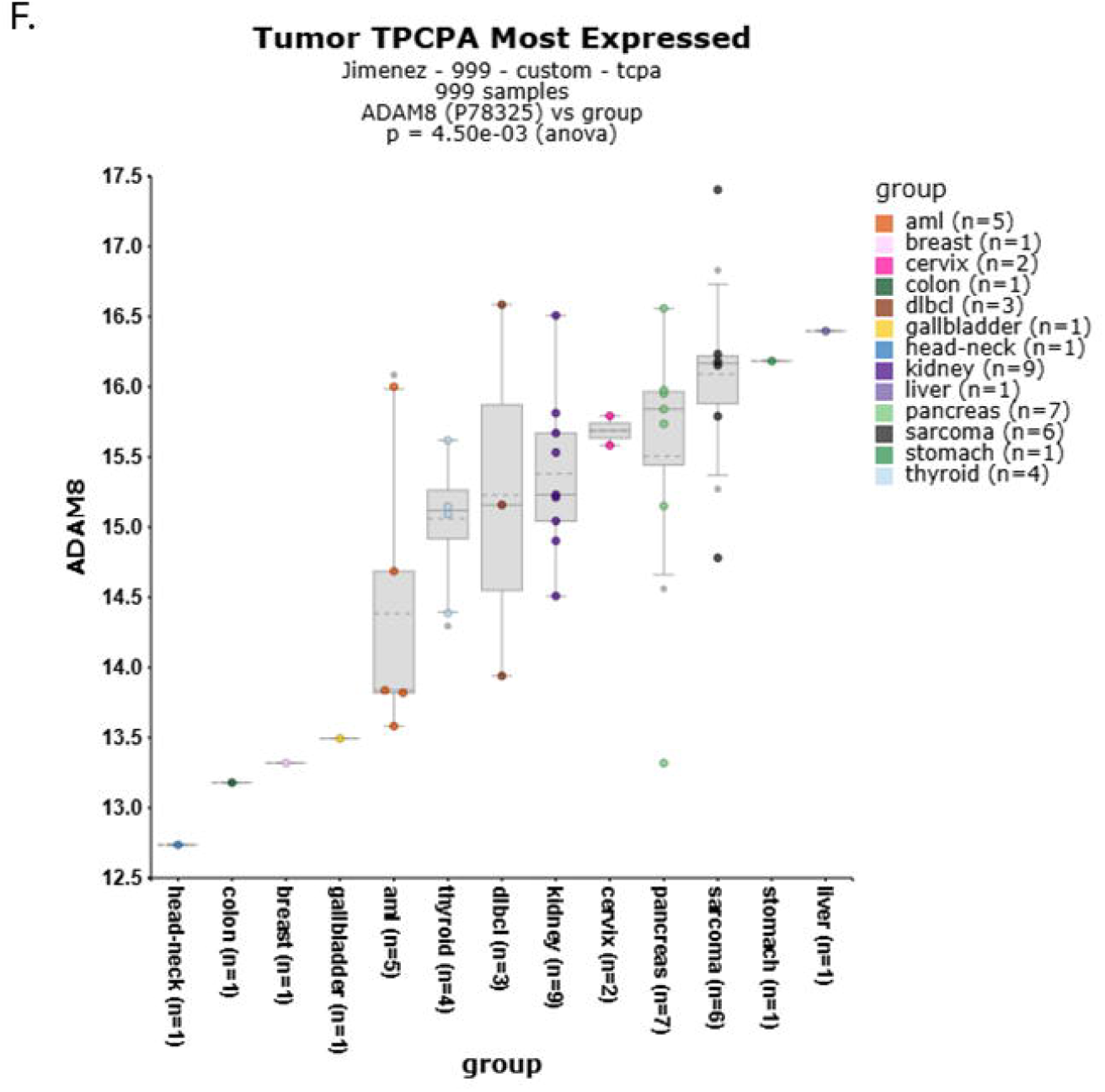
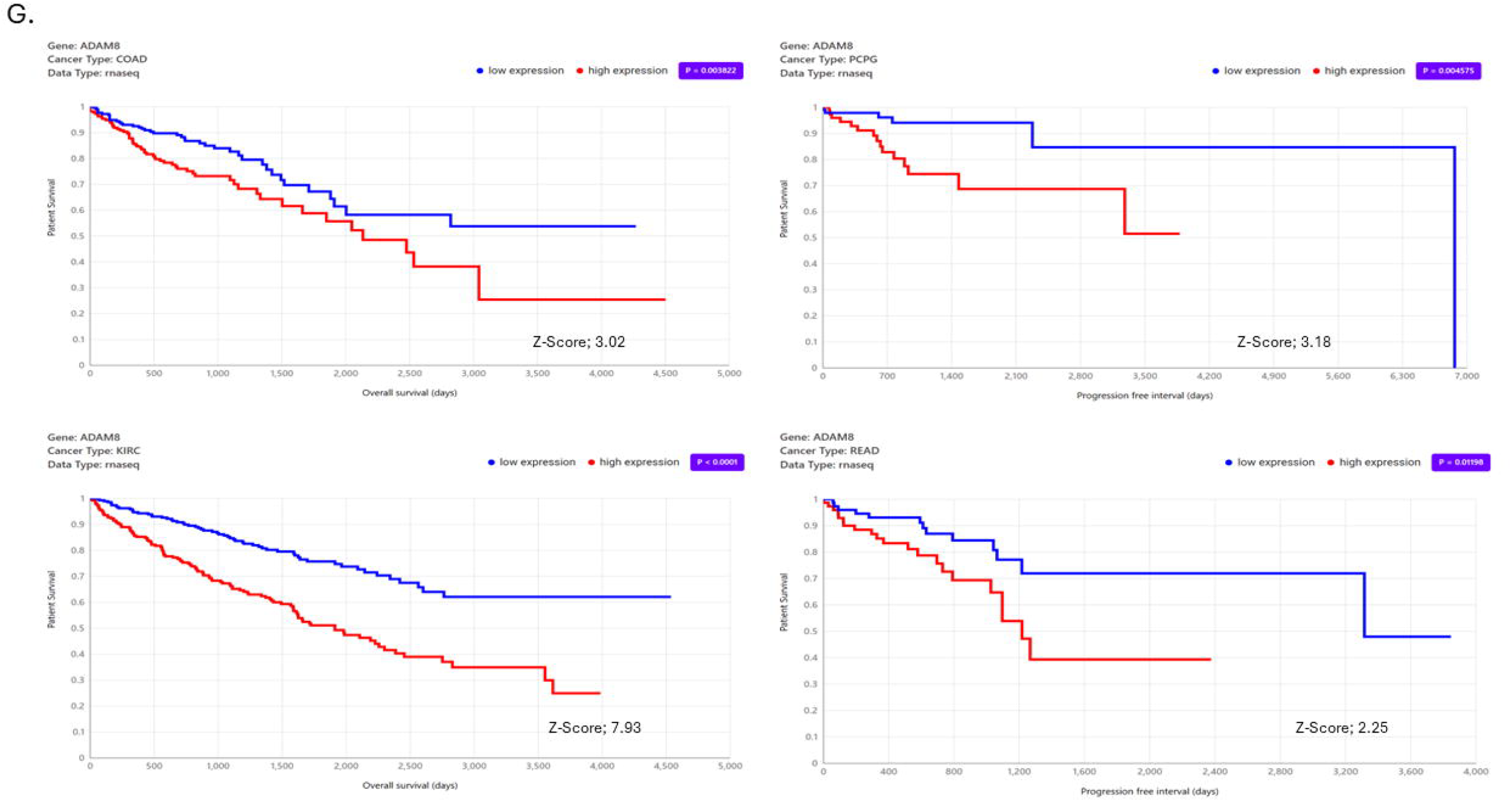
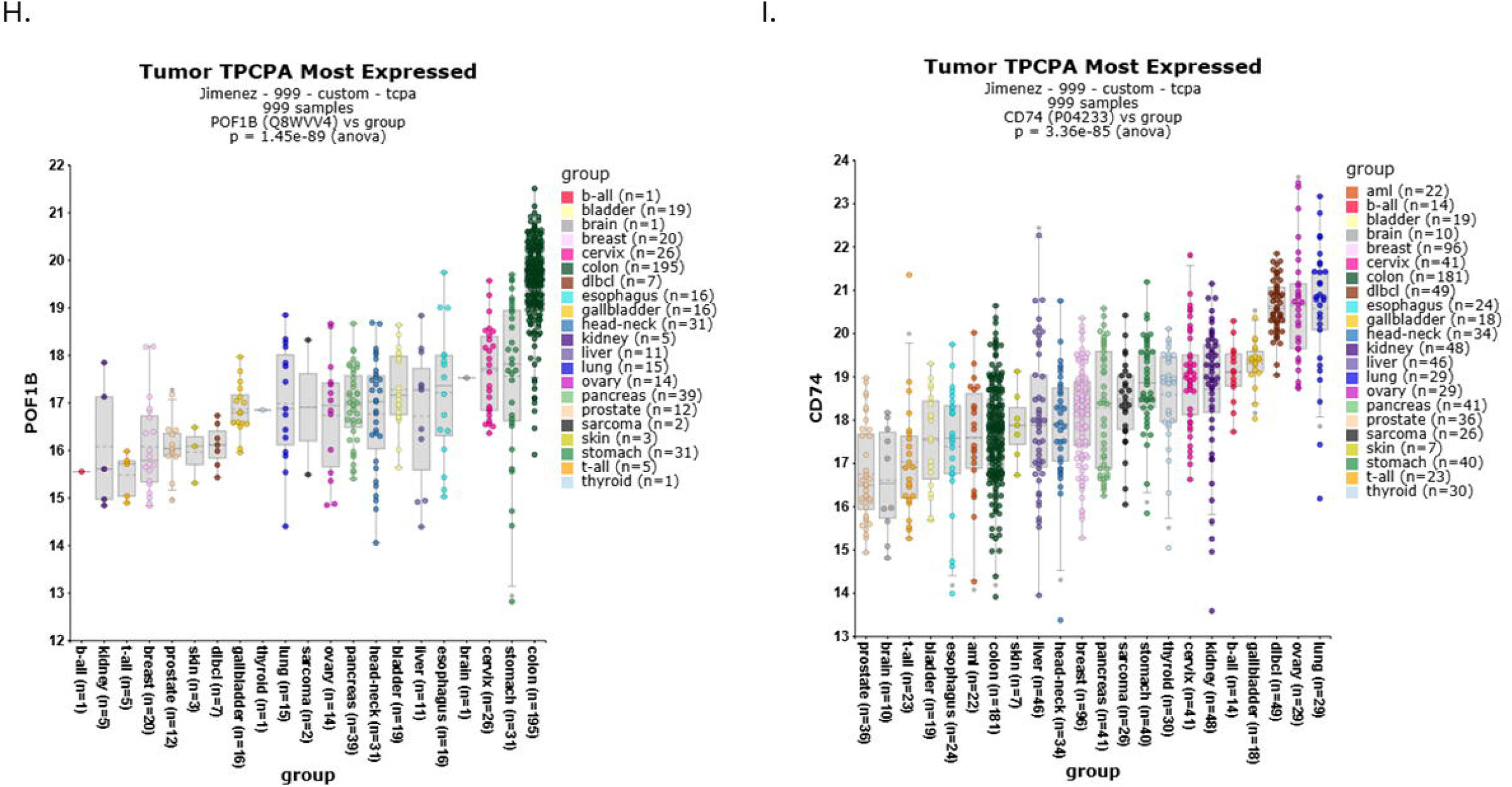
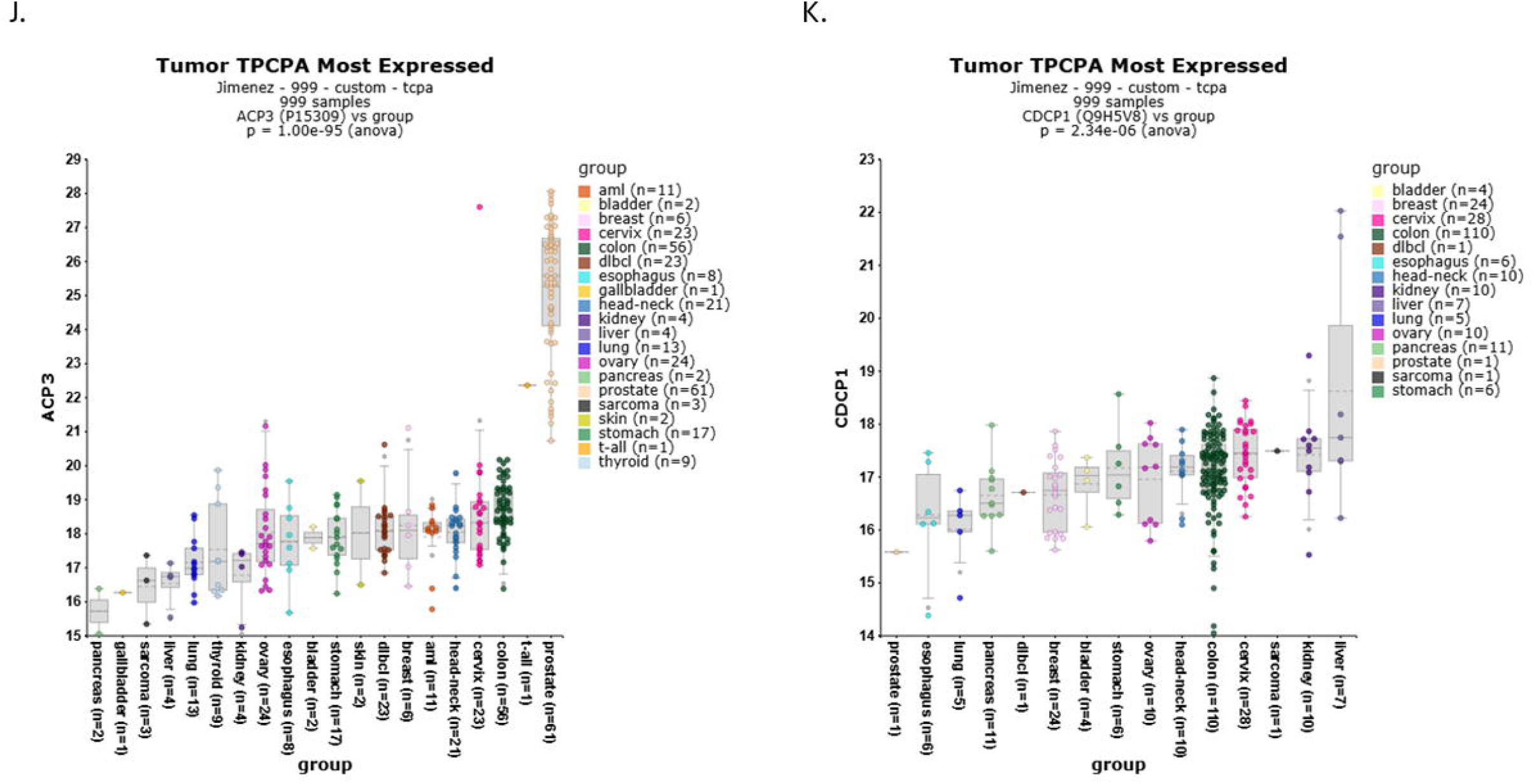
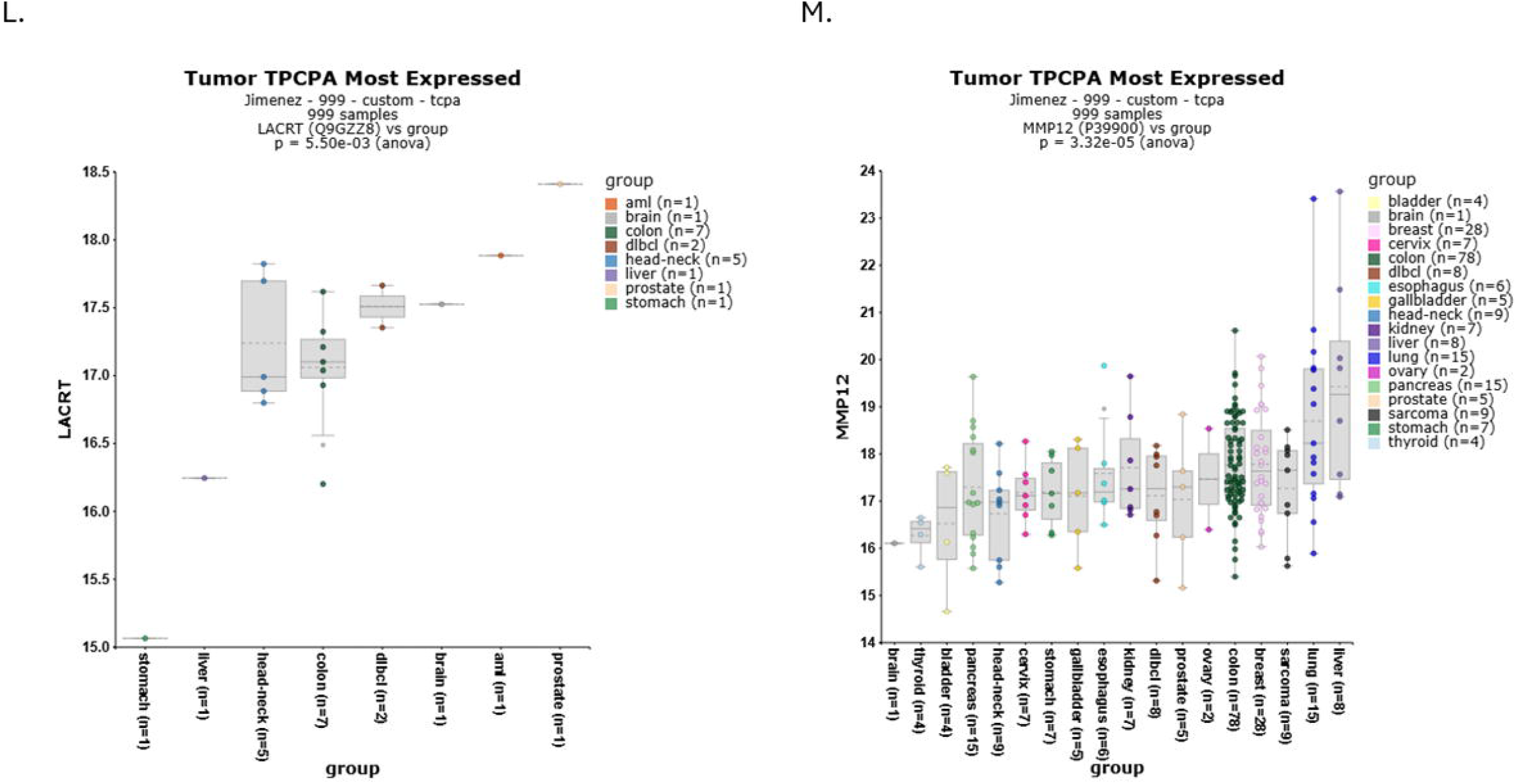
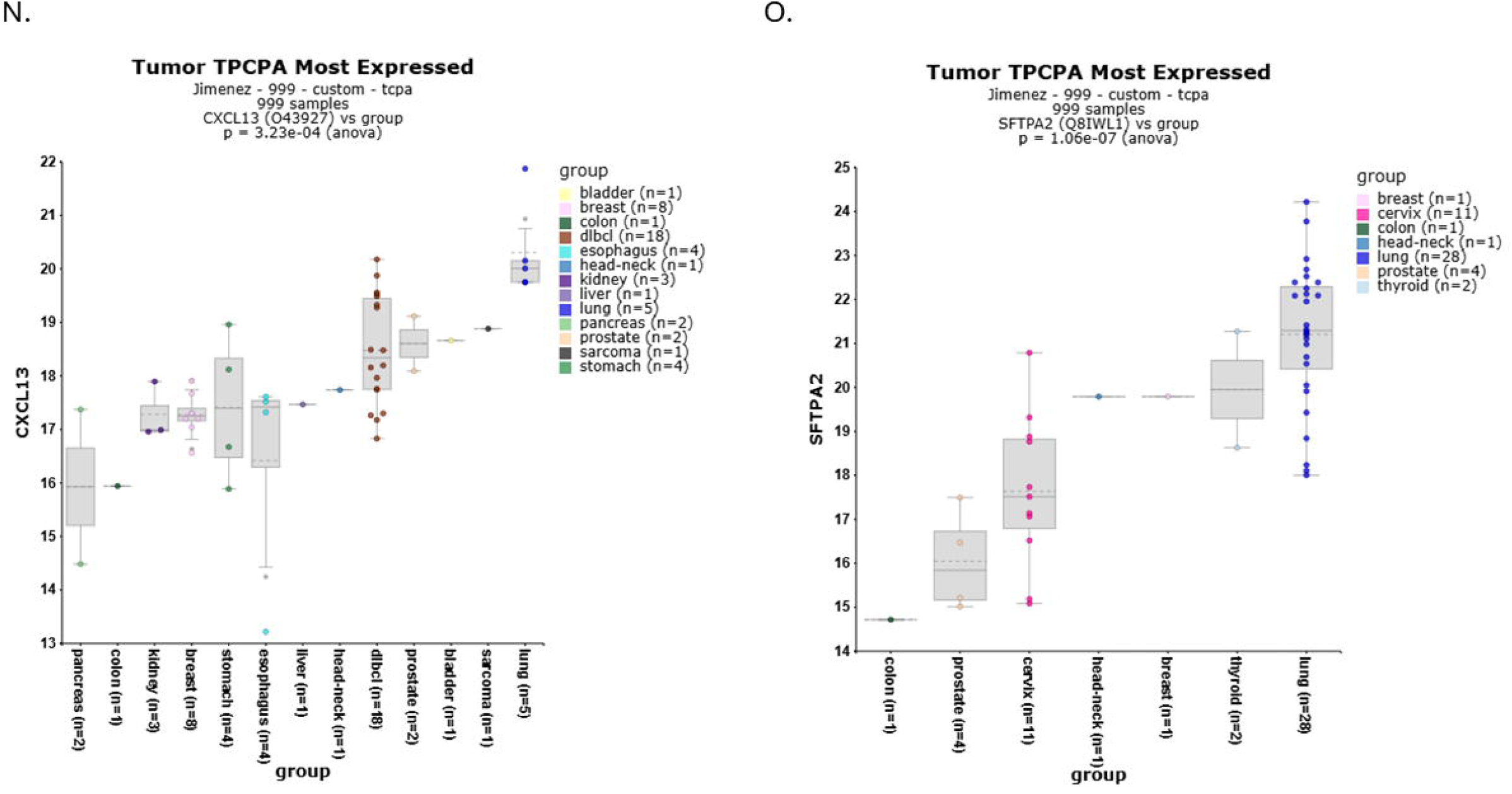
Cross-platform evidence for prognostic and proteomic signals across the candidate panel (GEPIA3.0 forests, TCGA-survival KM, and TPCPA proteomics). (A-C) GEPIA3.0 forest plots summarizing cancer-type–stratified Cox models for overall survival (OS) across the panel genes. Points show hazard ratios (HR) with 95% CIs; vertical line indicates HR=1. Multiple testing controlled by Benjamini–Hochberg (significant cohorts labeled). (D-E) Representative Kaplan–Meier curves from TCGA-survival for high vs low expression (median cutoff) illustrating adverse outcomes for high expression of secretory/TGF-β and protease markers (e.g., TGFB1, CDCP1, MMP12, ADAM8) in selected epithelial cancers; log-rank p-values and Z-scores shown where provided. (F-I) Kaplan–Meier examples for immune-context genes (CXCL13, CD74), demonstrating tumor-dependent effects (protective in inflamed/tertiary-lymphoid–rich contexts; neutral/adverse in immune-excluded settings). (J-M) TPCPA protein abundance across tumor types for secretory/matrix markers (e.g., TGFB1, CDCP1, MMP12, ADAM8). Distributions are shown as platform Z-scores (box/violin plots) with ANOVA FDR for group differences; higher-abundance cancers are annotated. (N-O) TPCPA protein abundance for immune-follicular/antigen-presentation markers (CXCL13, CD74), highlighting cancer-type–specific protein detectability consistent with transcript trends and microenvironmental composition. All panels use HGNC symbols, harmonized axis limits, consistent color palettes, and publication-resolution exports. “Cross-platform concordance” indicates agreement in direction across GEPIA forests (risk), TCGA-survival KMs (curve separation), and TPCPA (protein elevation).

## Discussion

In this systematic review, we found that blood-based proteomic biomarkers hold significant promise for the early detection of multiple cancers. However, their performance and clinical readiness vary across cancer types and study designs. Circulating protein signatures can indeed distinguish cancer patients from healthy individuals before symptom onset, as evidenced by both retrospective case–control studies and prospective cohort analyses. This discussion contextualizes our findings, explores implications for clinical implementation, and examines limitations that must be addressed.

### Convergence of evidence on key biomarkers

It is noteworthy that across independent studies and platforms, specific proteins emerge consistently. GDF15, for example, appears as a general marker of “ill health” that is elevated before diagnosis in a variety of cancers[5]. This consistency suggests that *systemic effects of tumors* (like cachexia or immune activation) might manifest in the blood years before diagnosis. However, such markers are not specific; GDF15 can rise in benign conditions as well. The inclusion of inflammatory cytokines and acute-phase proteins in many panels underscores that cancer often co-opts or triggers systemic inflammatory pathways. Conversely, truly cancer-specific proteins (e.g., tumor antigens shed from cancer cells) are rare in circulation, especially at early stages. One implication is that combinatorial markers are required: an optimal MCED proteomic test may include a mix of broad “sentinel” markers (to maximize sensitivity across cancers) and organ-specific markers (to guide localization and reduce false positives)[1]. Our review identified several candidates for each category – e.g., broad markers such as GDF15 or IL-6, and specific markers such as CA-125 (ovary), AFP (liver), PSA (prostate), etc. The challenge is balancing these to achieve high sensitivity without sacrificing specificity.

### Diagnostic performance and clinical implications

The high accuracies reported in some training-phase studies (AUCs of 0.95+ [1, 6]) must be interpreted cautiously. Experts warn that such results are likely overestimates due to overfitting and require rigorous external validation[1]. Indeed, the clinical utility of MCED tests remains unproven. As stated by commentators, it is “a very long way from being proven” whether these tests will reduce mortality or even be acceptable in screening workflows[1]. Our findings echo those concerns: while technical performance in controlled studies is promising, real-world screening performance will be lower. For instance, a model that achieved 90% sensitivity in a case–control setup might deliver only, say, 50% sensitivity when applied prospectively (due to lower tumor burden, spectrum effects, etc.).

Additionally, the positive predictive value (PPV) of any MCED test in an asymptomatic population is a critical metric. With cancer prevalence low (perhaps 1–2% in a screened cohort over a year), even 99% specificity yields some false positives. The PATHFINDER trial of a DNA-based MCED test reported a PPV of ∼43%[3] – meaning more than half of positive results were false alarms, albeit with follow-up protocols in place. Proteomic tests would need to match or exceed that specificity. The case–control proteomic studies achieved ∼99% specificity[1], but whether those levels hold in independent sets remains unproven. False positives not only cause anxiety and invasive follow-ups[3] but also could harm the credibility of a screening program. Therefore, any proteomic panel must be prospectively validated in large cohorts (thousands) to estimate its factual specificity and PPV confidently.

On the positive side, if a proteomic test modestly improves early cancer detection for a subset of lethal cancers, the clinical impact could be significant. For example, lung cancer caught at Stage I has a 5-year survival of>70% vs <20% if caught at Stage III–IV. Our review found that a proteomic signature could potentially flag a proportion of lung cancers before they’re typically found (AUC ∼0.7 pre-diagnosis) [2]. If implemented in high-risk groups (e.g., smokers), such a test might complement imaging by identifying those who should undergo immediate low-dose CT (LDCT), thus refining screening efficiency. Similarly, the dire prognosis of late-stage ovarian cancer could be improved if a blood test (e.g., CA-125, HE4, plus novel markers) detects some ovarian cancers at an early, treatable stage. The UKCTOCS trial with CA-125 showed a stage shift (more early cases found) but did not significantly reduce mortality, likely because sensitivity was insufficient and follow-up procedures were complex. A more sensitive multi-marker test might achieve the mortality reduction if coupled with effective early intervention.

### Multi-cancer vs. organ-specific screening

A key advantage of pan-cancer blood tests is the potential to screen for multiple cancers simultaneously, including those without current screening (ovarian, pancreatic, kidney, etc.). However, this breadth brings complexity. Different cancers have different prevalence and biomarker profiles. Our review indicates that a one-size-fits-all panel will likely overperform some cancers and underperform others. One strategy is a “multi-tier” screening approach: use a broad proteomic test to indicate *any cancer signal*, then use secondary tests targeted to likely cancer types (based on marker patterns) to hone in on the diagnosis [1]. For instance, a positive “cancer signal” with a strong CA-125 and WFDC2 component would point to pelvic imaging for ovarian cancer. In contrast, a positive signal dominated by CEA and CYFRA21-1 might prompt a colonoscopy or chest CT. Such an approach is being evaluated (e.g., in the ongoing NHS Galleri trial for the DNA test). Proteomic tests need similar management pathways. Another point is integration with risk factors. Proteomic data can be combined with clinical variables to improve overall prediction[5]. For example, incorporating age, genetic risk scores, or smoking history with protein markers yields more robust models[6]. The Lancet Digital Health study found that, across 52 diseases, a sparse protein model outperformed traditional risk factors [6]. This suggests that in practice, a future screening algorithm might use a risk calculator (with demographic and proteomic inputs) to determine an individual’s “cancer risk score”, rather than a binary test.

### Readiness for translation

At present, no proteomic multi-cancer test is FDA-approved or widely deployed. The most advanced MCED test in trials is a DNA-based test. Proteomic tests are catching up, with some (e.g., the Novelna 10-protein panel [1]) possibly entering validation. Key steps toward translation include:

- **Prospective clinical trials:** We need extensive studies in which an unbiased cohort undergoes the proteomic test and is followed to determine which cancers are detected and which are missed. This provides true sensitivity, specificity, PPV, and NPV in practice. Encouragingly, some trials are underway (for instance, the SUMMIT study in the UK is evaluating various blood markers in ∼100k individuals; and the PROMISE study in China is testing a multi-omics panel). Our review underscores that without such trials, performance claims remain speculative[1].
- **Regulatory and logistical considerations:** Proteomic assays (such as Olink) are currently run in specialized labs. For population screening, tests must be robust, automatable, and cost-effective. The Olink panels, while powerful, are expensive and require a cold chain, etc. Simplification to a targeted set of proteins (ideally measured by an affordable platform, e.g., immunoassay or mass spec in clinical labs) will be needed. Many groups are now trying to reduce their discovery panels into an affordable chip or kit.
- **Economic and ethical implications:** Multi-cancer screening will generate new complexities – how to counsel individuals with a “positive cancer signal” but no symptoms and an uncertain anatomic source? This could lead to a cascade of imaging and biopsies. False negatives are also an issue; a negative test might falsely reassure someone who actually harbors a cancer that the proteomic panel doesn’t detect well (e.g., an early breast cancer). Therefore, these tests likely would *complement* – not replace – existing screening. They might be most useful for cancers that we currently do not screen for. Health-economic modeling will be needed to determine if the benefits (earlier detection, lives saved) outweigh potential harms and costs.

Collectively, our pathway/hallmark synthesis (Figure 1) and cross-platform validation (Figure 2) delineate a coherent matrix–immune–secretory axis underpinning multi-cancer detectability and prognosis. “Known” markers primarily track epithelial antigens and host inflammatory tone, whereas the “Novel” panel captures ECM protease/remodeling (MMP12, ADAM8), antigen presentation/B-cell follicular programs (CD74, CXCL13), secretory/TGF-β signaling, and epithelial invasion (CDCP1). Concordant signals across GEPIA3.0 forests (adverse HRs for secretory/matrix genes; context-dependent effects for immune markers), TCGA-survival Kaplan Meier curves (significant separations in representative cohorts), and TPCPA proteomics (tumor-type–specific protein elevation) support a pragmatic two-bucket strategy: pair high-risk secretory/matrix markers with immune-context sensors to improve detection sensitivity, tissue-of-origin interpretability, and clinical triage. While platform antibodies, cohort heterogeneity, and portal defaults introduce limitations, the repeated agreement across transcript, survival, and protein layers argues for prospective translational testing, particularly in liquid-biopsy workflows where complementary panels can mitigate single-marker weaknesses and better reflect tumor–host ecology.

### Limitations of this review

Our meta-analysis has some limitations worth noting. First, the field is very new and evolving; many studies are heterogeneous in design, making formal quantitative synthesis challenging. We mitigated this by focusing on qualitative synthesis and highlighting representative studies rather than attempting to pool all data. Second, there may be publication bias – studies with exciting results (high AUCs) are more likely to be published. In contrast, negative or inconclusive findings (e.g., “proteins X, Y, Z did *not* predict cancer”) might be underreported. This could inflate our sense of how well proteomics works. Third, our inclusion was restricted to 2020 onward and to English-language studies, potentially excluding earlier critical work (such as the 2018 CancerSEEK study or non-English cohort studies). We referenced some historical benchmarks for context but did not systematically review pre-2020 studies. Finally, as an academic meta-analysis, we did not have individual patient data; thus, we relied on published summary statistics. Different studies might define metrics differently (one might report sensitivity at fixed specificity, another might report an optimized Youden’s index, etc.), which we had to interpret on common ground.

## Limitations

Several limitations are inherent to both the underlying studies and our review. Many of the primary studies were proof-of-concept studies with relatively small sample sizes, especially by cancer type [1]. This raises the risk of overfitting complex machine learning models can spuriously perform well on training data but fail in independent cohorts[1]. The lack of external validation in some studies (e.g., Budnik et al.’s 10-protein panel was not tested on an independent cohort) means we must view their impressive accuracy with caution [1]. Where validations were performed (as in Uhlén et al.’s study using UK Biobank data), performance often dropped for certain cancers, underscoring concerns about overfitting and the real-world complexity [2]. There is also the issue of confounding and reverse causality in prospective analyses. Some proteins might be elevated due to subclinical disease or comorbid conditions rather than causally associated with cancer. For example, an individual developing lung cancer might have an inflammatory condition that increases cytokine levels, so the protein is a marker of the general risk environment, not a tumor secretome per se. Genetic analyses (Mendelian randomization) in the UK Biobank study sought to address this, identifying a few proteins with likely causal roles (e.g., SFTPA2 in lung cancer)[5]. However, most associations remain associative. Our review does not differentiate between causal and reactive biomarkers, but this distinction matters if one aims to intervene on the pathway rather than use it as a marker. A limitation of proteomic technology is that current assays cover only a fraction of the proteome [5]. As noted, the Olink panels measured ∼1.5k to 5k proteins; there are >20k proteins in the human proteome, and potentially many more modified forms. It’s possible that some highly tumor-specific markers (e.g., cancer-testis antigens, neoantigens, or low-abundance signals) are not yet included in these panels. As technology advances to measure more analytes (or new types, such as post-translational modifications), the biomarker landscape could shift. Our review is thus a snapshot based on what’s measurable today. On the review side, we did not perform a formal risk-of-bias grading for each study (e.g., QUADAS-2 tool for diagnostic accuracy studies) due to the heterogeneity of study designs. Nonetheless, we qualitatively commented on biases (e.g., selection of convenience cohorts). We also acknowledge that our summary table, while covering key studies, is not exhaustive. There are likely ongoing trials and unpublished data (especially from industry) that we could not include. Finally, our review focused solely on *proteins*, but many emerging MCED approaches are multi-omics. We included some references to multi-omics (e.g., the PROMISE study), in which proteins were a component, but we did not comprehensively review DNA or other markers. Some might view this as a limitation, but it was a deliberate focus to delve deeply into proteomics. In practice, the synergy between proteins and DNA markers could be crucial—for example, a modest protein panel combined with a ctDNA test might yield much better performance than either alone [1]. This combined performance was outside our scope but is an essential consideration for the future of MCED.

## Conclusion

Blood-based proteomic biomarkers have rapidly advanced as candidates for multi-cancer early detection, leveraging the ability to capture tumor and host responses in a single assay. Our comprehensive review indicates that while no single protein is a silver bullet for cancer detection, multi-protein signatures can achieve notable accuracy in distinguishing cancer from healthy states across a range of cancer types. In particular, extensive prospective studies have shown that specific proteins begin to deviate from normal levels years before a cancer diagnosis – offering a window for potential early intervention [2, 7]. Promising panels, some comprising fewer than a dozen proteins, have shown extremely high sensitivity and specificity in preliminary studies[1]. Though these results await external validation, they suggest that proteomics could complement, or even rival, genomic assays in MCED applications. However, translating these findings into clinical screening programs will require overcoming several challenges. First, extensive validation in diverse, prospective cohorts is essential to confirm performance and generalizability. Ongoing trials and population studies (e.g., in the UK, the US, and China) will be informative in the next 2–3 years. Second, optimization of assay platforms for cost, speed, and scalability is needed. Current high-throughput proteomics may be impractical for millions of tests, so simplified targeted kits must be developed containing the most informative markers. Third, clinical pathways for managing test results need to be defined: for instance, what diagnostic work-up follows a “positive” proteomic test in an asymptomatic person? As MCED tests become more real, healthcare systems must prepare for those downstream implications (including potential overdiagnosis). Our review highlights that certain cancers (e.g., lung, ovarian, liver) appear ripe for proteomic screening, given strong blood signals and the lack of existing screening tools. Others, like breast and prostate, may require additional strategies or combined modalities to achieve acceptable detection rates. It is conceivable that future screening could employ a panel of panels e.g., a core proteomic test plus cancer-type-specific reflex tests (which might be other proteins or imaging). The high specificity required for population screening also suggests that a multi-step approach (an initial broad test followed by a confirmatory test) could optimize the balance between detecting cancer and minimizing false alarms [3]. In conclusion, circulating proteomic signatures represent a frontier in early cancer detection, now being illuminated by large-scale research. The evidence to date is encouraging that proteins in the blood can reveal the hidden presence of cancer, sometimes well before clinical manifestation [2]. Several candidate protein panels have demonstrated the ability to detect multiple cancers with accuracy that, if confirmed, would have a profound clinical impact—enabling a single blood test to screen for many cancers that today go undetected until advanced stages. Nonetheless, we must temper optimism with rigorous validation – the “hype” vs. “hope” dichotomy will be resolved only through careful clinical evaluation [1]. As one expert noted, “if the assay performance in future studies is anywhere close to what this preliminary study suggests, then it could really be a game changer”[1]. The coming years will determine whether proteomics-driven MCED tests can fulfill this promise. For now, they stand as one of the most intriguing developments in cancer prevention research, meriting continued large-scale investigation and cautious optimism for their role in saving lives through earlier diagnosis.

## Author information

### Authors and Affiliations

Department of Biochemistry, King George’s Medical University, Lucknow, Uttar Pradesh, India 226003.

Department of Pathology, King George’s Medical University, Lucknow, Uttar Pradesh, India 226003.

Vivek Singh & Rashmi Kushwaha

### Contributions

VS wrote the manuscript and made figures. RK analyzed the manuscript. The author(s) read and approved the final manuscript.

## Ethics declarations

### Ethics approval and consent to participate

Not Applicable.

### Consent for publication

Not applicable.

